# Multi-omics characterization of the skin microbiota reveals the anti-aging roles of *Stenotrophomonas maltophilia*

**DOI:** 10.1101/2025.07.09.663908

**Authors:** Danni Guo, Yuanyuan Chen, Yahong Wu, Jingmin Cheng, Wenjie Lai, Wentao Ma, Hang Yang, Lianyi Han, Lan Ma, Haidong Jia, Xiao Liu

## Abstract

Shifts in the skin microbiome have shown a close link to chronological age. However, the contribution of skin microbiome in skin aging phenotypes remains unclear. To explore this, we performed phenotypic, metabolomic, metagenomic, and functional analyses on a cohort with divergent skin aging phenotypes. Genome-scale metabolic models (GEMs) integrated with metabolomic analysis revealed that *Stenotrophomonas maltophilia*, enriched in the younger group (categorized by AI-predicted age and skin elasticity), utilizes the glutathione cycle to maintain redox homeostasis. Cellular experiments showed its metabolites enhanced GSH synthesis and alleviated oxidative stress-induced skin aging by upregulating key genes in fibroblasts, including *GCLM*, *PGD*, *SOD2*, and *NQO1*. Additionally, GEMs highlighted its potential anti-aging roles in regulating host metabolic pathways involving betaine, lysolecithin, and porphyrin. In parallel, *Acinetobacter guillouiae* was found to influence host melanin metabolism by degrading dopamine (DA) and 3-methoxytyramine (3-MT), offering potential therapeutic strategies for mitigating pigmentation. Our findings highlight the dynamic interplay between skin microbiota and the host in aging, offering new insights for designing targeted anti-aging interventions.

## Background

The skin is the largest organ of the human body^1^. Beyond the gastrointestinal tract, it also harbors the second-most extensive microbial communities in the human host^2^. The human skin microbiome is a complex ecological system comprising bacteria, archaea, fungi, and viruses^3^. This microbiome interacts reciprocally with the host’s skin health, influencing conditions ranging from its general health to specific diseases and its aging state^4–6^. Resident skin microbes help maintain skin homeostasis and can modulate inflammatory responses or induce pathogenicity based on host environmental conditions^7–9^.

Skin aging is a multifaceted biological process influenced by a myriad of intrinsic and extrinsic factors. Typically, with age progression, a gradual decline in cellular regeneration occurs, accompanied by diminished DNA repair capacities. Concurrently, there is a deterioration in the fundamental supporting structures of the skin, accelerating the process of skin aging^10^. Additionally, environmental determinants, notably prolonged exposure to ultraviolet (UV) radiation, can amplify the generation of reactive oxygen species within cellular structures, thereby causing oxidative stress. This heightened stress damages cell membranes, destabilizing the skin’s integral collagen and elastin fibers, and compromising DNA integrity, consequently accelerating skin cell aging^11^.

Physiological alterations brought about by skin aging manifest as increased wrinkling, loss of elasticity, and a decrease in water content in the stratum corneum, and reduced sebum production^12–15^. Skin aging also represents an ecological shift in the skin, impacting its resident microbial community. Several studies have unveiled a complex interplay between skin aging and the skin microbiome, including changes in microbial diversity and composition as age progresses^16–19^. These alterations extend beyond mere changes in microbial types and quantities, encompassing modifications in their metabolic pathways and functions. Importantly, variations within these microbial communities have been closely linked to critical indicators of skin health, including pigmentation, collagen synthesis, sebum production, and the moisturizing capability^20–22^. Specifically, changes in microbial abundance and metabolic activity influence these key physiological processes, thereby accelerating or decelerating the external manifestations of skin aging^23^.

With growing research into the intricate dynamics of skin aging and its microbial influences, some studies have proposed that the phenotypes of skin aging and susceptibility might more accurately reflect the characteristics and changes in the skin microbiome than chronological age itself^24–26^. However, most of these studies relied on population-based cohorts with varied groups of chronological ages, in which ages may serve as a confounding factor. There is growing evidence that the age of 40s are considered the watershed of skin aging^27–29^. During this period, collagen production decreases, and menopause accelerates this by lowering estrogen levels. Slower cell turnover and increased oxidative stress make the skin more vulnerable to free radical damage, speeding up the formation of wrinkles and dryness. Despite this recognized importance, studies specifically focusing on the relationship between microbial communities and skin aging phenotypes within this pivotal age group remain scarce.

Thus, we conducted a comparative study on 103 healthy women at the age of 39-41 exhibiting divergent conditions of skin aging. Using metagenomic analysis, we identified skin bacterial markers associated with aging conditions. Additionally, by constructing GEMs and implementing non-targeted metabolomics, we have elucidated the skin aging-related metabolic networks co-regulated by the skin microbiome and the host. Specifically, we identified and functionally validated the anti-aging role of skin commensal *Stenotrophomonas maltophilia*. Our comprehensive analysis elucidates the microbial contributions to the skin aging trajectory. The microbial markers and metabolites we identified hold substantial potential for aiding the development of anti-aging therapies.

## Results

### Diverse phenotypes across skin aging groups

In this study, 202 healthy Chinese women aged 39 to 41 were enrolled, and we utilized the artificial intelligence platform Megvii’s Face^++^ to classify participants into two age groups^30^: 100 individuals with AI-estimated ages significantly below their chronological age (younger group) and another 102 with AI ages notably above (older group). Key skin traits were assessed across both groups. Using elasticity R2 as a principal metric, we selected the top 52 younger participants with the highest elasticity scores and the 51 older participants with the lowest scores for further analysis. We assessed skin physiological traits and examined skin microbiome and metabolites separately using shotgun metagenomics and non-targeted LC-MS (**Fig. 1a**). Skin phenome analysis revealed that the younger group exhibited significant enhancements in skin elasticity index R2, sebum content, porphyrin optical density, porphyrin area percentage, and stratum corneum hydration, suggesting better moisturization and overall skin health compared to the older cohort (**Fig. 1b, Table S1**). As anticipated, the older group exhibited more wrinkles. There were also notable differences in skin pigmentation between the groups, with the older group showing a significant reduction in Luminance (L*) and marked increases in redness (a*) and yellowish tones (b*), suggesting greater pigment accumulation with skin aging. Additionally, the lack of significant differences in Transepidermal Water Loss (TEWL) and pH levels between younger and older groups was observed (**Fig. 1b**). Correlation analysis revealed that most skin phenotypes were strongly correlated with AI-estimated age, confirming its efficacy as a reliable indicator of skin aging. Moreover, wrinkles appear to be an integrative manifestation of aging, reflecting diminished stratum corneum hydration and sebum levels, alongside increased pigment deposition which may be associated with elevated oxidative stress possibly due to sun damage^31^. These findings highlight the complex interactions and regulatory mechanisms between various skin aging indicators (**Fig. 1c**).

**Fig. 1.**
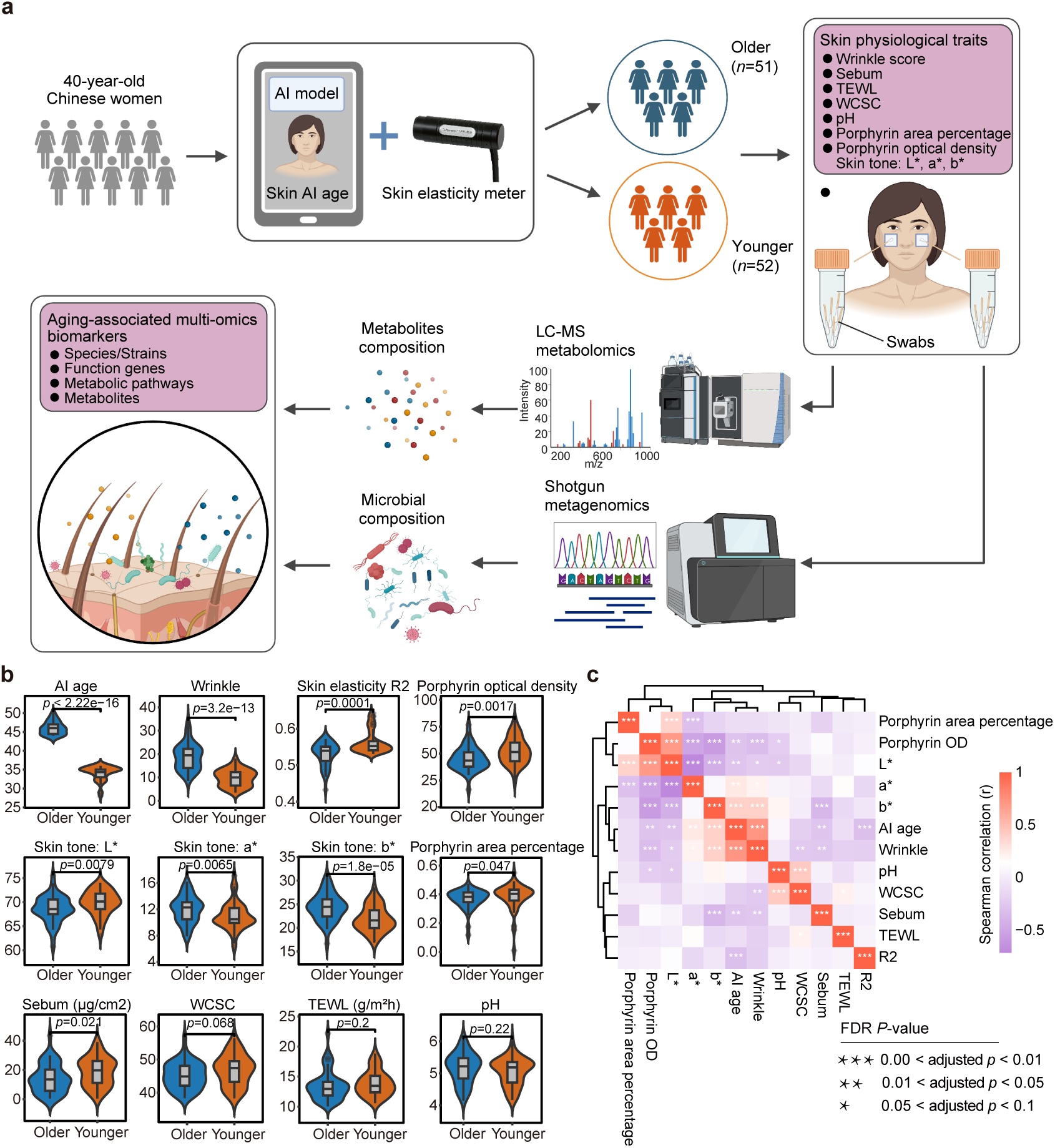
Overview of the study design and comparative skin aging phenotypes in women of the same age group. (a) The overall workflow of this study. (b) Violin plots show the substantial differences in skin phenotypes between the younger and older groups, analyzed using the Wilcoxon test. AI age: Estimated age using artificial intelligence; Skin elasticity R2: Skin elasticity measured by the R2 parameter; Skin tone: L*: Skin lightness; a*: Skin red-green component; b*: Skin yellow-blue component; Porphyrin area percentage: Percentage of skin area covered by porphyrin; WCSC: Water Content of the Stratum Corneum; TEWL (g/m²·h): Transepidermal Water Loss. (c) Heatmap of Spearman’s correlation among skin aging phenotypes. Purple hues indicate negative correlations, with the color intensity corresponding to the strength of the correlation. Significance levels are denoted as follows: *, adjusted *P* < 0.1; **, adjusted *P* < 0.05; ***, adjusted *P* < 0.01.

### Skin metagenomics reveals aging-associated dynamics in the skin microbiome

From the metagenomic analysis, we identified 2,419 microbiome species, with 1,464 species shared across the younger and older groups (**Figure S1a, Table S2a**). Among the top 20 species, *Cutibacterium acnes* (*C. acnes*) was the most abundant resident skin microorganism, followed by *Staphylococcus epidermidis* and *Staphylococcus hominis* (*S. hominis*) (**Fig. 2a**). Our data revealed significantly lower Shannon diversity of microbial species in the older group (**Fig. 2b, Table S2b**). Previous studies, primarily focusing on chronological age, suggested that older individuals had higher microbial diversity in their skin^17,18,22,23,32^. Notably, it has also been reported that a decrease in *C. acnes* abundance in older individuals might lead to an increase in species diversity^19,20^. However, our samples, derived from women of the same age range, showed no significant difference in *C. acnes* abundance between groups (adjusted *P* = 0.11, Log_2_FoldChange = 0.39, **Table S2d**). We hypothesize that the lower microbial diversity observed in younger groups in prior studies might be due to increased *C. acnes* abundance. To verify our hypothesis, we analyzed a dataset of 742 samples, which previously categorized C-cutotypes as representative of young groups and M-cutotypes of old groups^32^. When removing the influence of *C. acnes* abundance on the microbial diversity between groups, the data also supported a higher microbial diversity in the young group (**Figure S1b, Table S2e**). Thus, our results provide additional evidence than those of earlier studies that a higher level of microbial diversity is associated with both greater ecological stability and younger skin.

**Fig. 2.**
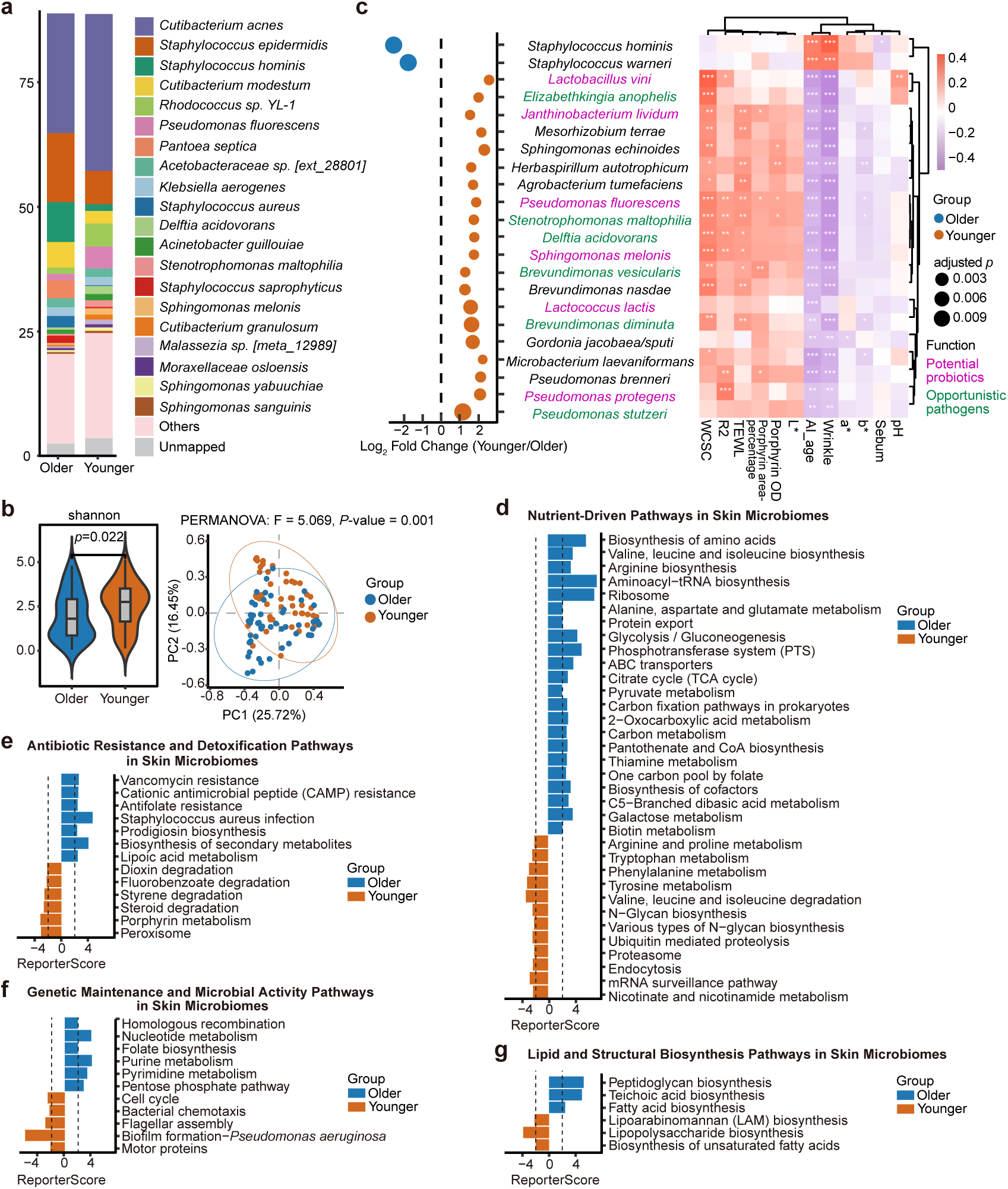
Comprehensive analysis of skin metagenomics. (a) Top 20 microbial species by relative abundance identified using mOTUs3. (b) Violin plot illustrating the Shannon index across groups, highlighting differences in microbial diversity, analyzed using the Wilcoxon test. Accompanying PCoA of Bray-Curtis distances plot depict β-diversity among microbial species, emphasizing group variations. (c) Left panel: Bubble chart displaying microbial species with significant differences between groups (absolute Log_2_FoldChange > 1, adjusted *P* < 0.05). Right panel: Heatmap showing the Spearman correlation analysis between differential microbial species and skin phenotypes. Significance levels are indicated as follows: *, adjusted *P* < 0.1; **, adjusted *P* < 0.05; ***, adjusted *P* < 0.01. (d) Nutrient-driven pathways enriched in skin microbiomes, with a cutoff Reporter score > 1.96. The older group shows significant enrichment in amino acid biosynthesis, protein synthesis, and carbon and energy metabolism pathways, suggesting an adaptive response to nutrient scarcity in aging skin. (e) Antibiotic resistance and detoxification pathways enriched in skin microbiomes, with a cutoff Reporter score > 1.96. The older group exhibits increased resistance to antibiotics and greater susceptibility to infections, while the younger group is enriched in pathways for detoxifying environmental pollutants. (e) Genetic maintenance and microbial activity pathways enriched in skin microbiomes, with a cutoff Reporter score > 1.96. The older group shows enrichment in nucleotide metabolism, folate biosynthesis, purine metabolism, pyrimidine metabolism, and homologous recombination pathways, suggesting higher activity in genetic maintenance. The younger group is enriched in pathways related to cell cycle, bacterial chemotaxis, flagellar assembly, and motor proteins, indicative of higher cellular interaction and mobility. (f) Lipid and structural biosynthesis pathways enriched in skin microbiomes, with a cutoff Reporter score > 1.96. The younger group is enriched in pathways for maintaining membrane fluidity and immune modulation, while the older group is enriched in pathways for maintaining cell wall integrity and structural components.

β-diversity analysis indicated significant differences in microbial species composition between groups (**Fig. 2b**). Differential analysis identified 43 microbial species uniquely enriched across groups (**Figure S1c, Table S2d**). Notably, *S. hominis*, which was more prevalent in the older group, exhibited a significant negative correlation with sebum content. This observation aligns with the conclusions of previously published studies^33^. The decline in sebum content with aging may create a favorable ecological niche for *S. hominis* (**Fig. 2c**). Moreover, potential probiotics and opportunistic pathogens were more prevalent in the younger group (**Fig. 2c**). For instance, *Lactococcus lactis and Janthinobacterium lividum* were enriched, known for lipid regulation and antifungal activity^34–38^. Notably, *S. maltophilia*, though typically opportunistic^39–41^, correlated positively with youthful traits such as hydration and elasticity (**Fig. 2c**), suggesting some pathogens may support skin youthfulness under healthy conditions.

Our findings reveal an aging-related metabolic shift in the skin microbiome (**Fig. 2d-g, Table S2g**). In older skin microbiomes, enrichment of amino acid and aminoacyl-tRNA biosynthesis pathways (**Fig. 2d**) suggests adaptation to nutrient-limited conditions via increased reliance on internal synthesis to support protein and energy needs^42,43^. Conversely, younger skin microbiomes exhibit enhanced amino acid metabolism (**Fig. 2d**), particularly in arginine and proline pathways, supporting immune modulation, oxidative stress resistance, and environmental resilience^44–50^. Additionally, the older skin microbiome showed a notable accumulation of antibiotic resistance genes, including those resistant to vancomycin, cationic antimicrobial peptides (CAMP), and antifolates (**Fig. 2e**), suggesting adaptation to host immune pressures^51^ and potentially contributing to *Staphylococcus aureus* susceptibility in aged skin^52,53^ (**Fig. 2e**). Conversely, younger skin microbiomes play crucial roles in detoxifying environmental pollutants, such as dioxins^54–56^, fluorobenzoates^57,58^, and styrenes^59^, and in metabolizing harmful chemicals through pathways like porphyrin metabolism (**Fig. 2e**). Moreover, older skin microbiomes exhibited an enrichment in nucleotide synthesis and degradation pathways, such as purine metabolism and folate biosynthesis, suggesting a higher activity in DNA and RNA synthesis and repair (**Fig. 2f**). In contrast, pathways like cell cycle, bacterial chemotaxis, flagellar assembly, and motor proteins were primarily enriched in younger skin microbiomes, indicative of higher cellular proliferation and mobility (**Fig. 2f**). These patterns highlight a shift from growth-oriented activity in younger skin to genomic maintenance in older skin. Furthermore, lipid and structural biosynthesis pathways showed aging-related differences (**Fig. 2g**). Younger microbiomes favored unsaturated fatty acid and lipopolysaccharide biosynthesis, supporting membrane function, while older microbiomes showed enrichment in fatty acid and cell wall biosynthesis, reflecting structural maintenance. Overall, the enriched microbial pathways reflect adaptations to the shifting skin microenvironment with age, likely linked to changes in cellular turnover, nutrient availability, and immune function.

### Skin metabolomics sheds characteristic insights on potential targets of skin aging

Metabolomic profiling of skin revealed distinct metabolic patterns between age groups. The older group was enriched in metabolites linked to oxidative stress, chronic inflammation, and the accumulation of potentially toxic compounds. In contrast, the younger group showed higher levels of metabolites with moisturizing, anti-inflammatory, and anti-melanogenic functions (**Fig. 3a**).

**Fig. 3.**
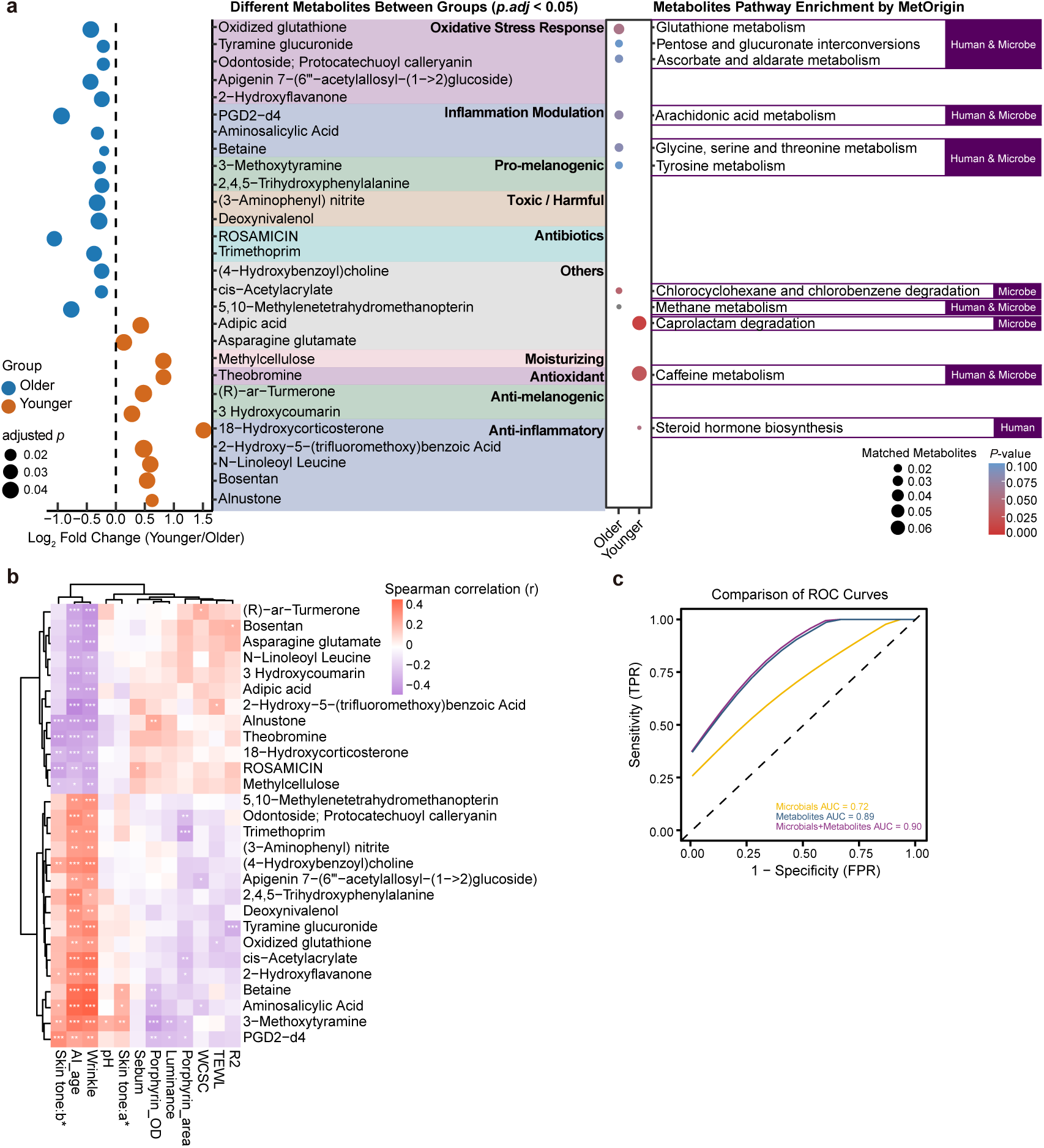
Comprehensive analysis of skin metabolomics. (a) Left: Bubble plot showing significantly different skin metabolites between the younger and older groups (adjusted *P* < 0.05). Bubble size reflects adjusted *P* values. Right: Enriched metabolic pathways identified using MetOrigin based on differential metabolites. The color indicates *P* value of pathway enrichment, and bubble size represents the ratio of matched metabolites. Annotations of pathway origin (Human, Microbe, Human & Microbe) indicate predicted potential sources of metabolites according to the MetOrigin database. (b) Heatmap illustrating the Spearman correlation analysis between differential skin metabolites and various skin phenotypes. Significance levels are indicated as follows: *, adjusted *P* < 0.1; **, adjusted *P* < 0.05; ***, adjusted *P* < 0.01. (c) Receiver Operating Characteristic (ROC) curves compare the predictive performance of models built using microbial data (orange), metabolomic data (blue), and combined microbial-metabolomic data (purple). The area under the curve (AUC) values are shown for each model: microbial data (AUC = 0.72), metabolomic data (AUC = 0.89), and combined data (AUC = 0.90).

In older skin, Glutathione metabolism and Arachidonic acid metabolism were significantly activated, with increased levels of oxidized glutathione and the inflammatory mediator PGD2-d4 (**Fig. 3a**), indicating a sustained oxidative and inflammatory state^31,60,61^. Additionally, consistent with previous findings of antibiotic resistance gene enrichment in the aged skin microbiome, we observed elevated levels of antibiotic-associated metabolites such as rosamicin and trimethoprim (**Fig. 3a**), likely reflecting microbial imbalance. The accumulation of fungal toxins such as deoxynivalenol may further impair lipid synthesis and membrane integrity, compromising skin barrier function^62^ (**Fig. 3a**). These combined alterations may contribute to skin aging.

In pigmentation-related metabolism, the older group showed elevated levels of 3-methoxytyramine (**Fig. 3a**), a product of Tyrosine metabolism and implicated in melanin biosynthesis. Skin phenotype correlation analysis indicated its positive correlation with skin tone parameters a* and b*, and negative correlation with L*, highlighting its significant role in pigment deposition in aging skin **(Fig. 3b).** In contrast, the younger group was enriched in (R)-ar-turmerone and 3-hydroxycoumarin (**Fig. 3a**), known tyrosinase inhibitors with photoprotective potential, which may serve as functional ingredients for pigmentation control^63–65^.

Additionally, we identified two metabolites not previously associated with skin function but significantly enriched in the younger group. N-linoleoyl leucine, formed through an amide bond between linoleic acid and leucine, was notably elevated (**Fig. 3a**). As linoleic acid plays a key role in immune regulation, anti-inflammation, and maintaining skin barrier integrity^66–68^, its derivative may similarly support skin protection through multiple pathways and represents a promising candidate for further investigation. We also observed enrichment of 2-hydroxy-5-(trifluoromethoxy)benzoic acid (**Fig. 3a**), a compound structurally similar to β-LHA (2-hydroxy-5-octanoyl benzoic acid), which is widely used for skin moisturization and anti-inflammatory applications^69–71^. While both share hydroxy and carboxylic acid groups, the trifluoromethoxy substitution in this compound enhances lipophilicity, chemical stability, and dermal permeability, potentially enabling deeper skin penetration and targeted anti-inflammatory, antibacterial, or antioxidant activity.

### Predictive modeling and construction of genome-scale metabolic models (GEMs)

To assess whether skin commensals or metabolites can classify premature or delayed skin aging, we built models using differential microbiome, metabolite, or combined features. Key features were selected using MeanDecreaseAccuracy, yielding 46, 205, and 133 features for the microbiome, metabolome, and combined models, respectively (**Table S3e**). The metabolomics model achieved strong predictive performance (AUC = 0.89), underscoring the importance of metabolic pathways in skin aging (**Fig. 3c, Table S3f**). In contrast, the microbiome model showed limited performance (AUC = 0.72; specificity = 93.3%) (**Fig. 3c**), suggesting that microbial data alone may not capture the complexity of aging. Combining microbiome and metabolomic features improved prediction (AUC = 0.90) (**Fig. 3c**), likely by capturing cross-domain interactions. To further explore this interplay, we constructed GEMs to investigate the functional roles of microbially derived metabolites in skin aging.

We constructed GEMs for 56 strain-level metagenome-assembled genomes (MAGs) with ANI >97.5%, covering 50 species (**Figure S3a, Table S4a**). Phylogenetic trees were constructed for different strains within the same species and were compared against previously annotated strains (**Figure S3b**). These MAGs accounted for ∼78% of total clean reads with no significant differences between age groups (**Figure S3c, Table S4b**), representing a substantial portion of the skin microbiome. MAG analysis revealed higher microbial diversity in the younger group, with *S. maltophilia* enriched and associated with youthful skin traits, while *S. hominis* was more abundant in the older group and negatively correlated with sebum production (**Figure**. **S3c–d, Table S4c,e**). These patterns are consistent with previous findings^33^.

GEMs were reconstructed for each MAG, comprising on average 2,028 reactions and 1,727 metabolites (**Table S5a**). Flux Balance Analysis (FBA) conducted to test model reliability demonstrated good stoichiometric and flux consistency, with no abnormally high ATP production under either aerobic or anaerobic conditions, confirming the models’ reliability (**Figure S4a**). Further, we weighted different MAGs by their relative abundance in samples and combined the GEMs of different samples from both older and younger groups to create a group-wide GEM. This combined model contained 4,739 reactions and 3,467 metabolites, with a flux consistency of 0.99 and stoichiometric consistency of 0.58, further confirming the reliability of the models (**Table S5a**). These validated models were subsequently used to investigate functional metabolic capacities of specific microbes and to explore metabolic differences associated with skin aging.

### *S. maltophilia* alleviates oxidative stress-induced skin aging

Oxidative stress is a pivotal factor in skin aging^31^. Free radicals and reactive oxygen species (ROS) damage skin cells, leading to the breakdown of collagen and elastin, thereby accelerating aging^72^. Previous studies have shown that glutathione (GSH), crucial for the cellular antioxidant defense system, neutralizes these free radicals and peroxides using the ascorbate-glutathione cycle to maintain cellular redox balance, converting into oxidized glutathione (GSSG) which protects cells from oxidative harm^73,74^ (**Figure S4c,d**).

Building on the findings of our study, which identified a high accumulation of GSSG in the skin of older individuals (**Figure S4b**), we focused on exploring the oxidative and reductive reactions of GSH within the skin microbiome. FBA analysis of the GEM revealed that the flux of GSH redox reactions was −2.36 in the younger group and −1.09 in the older group (**Table S5c**). These negative values indicate ongoing reduction processes of GSSG to GSH. The lower absolute value observed in the older group suggests reduced flux, consistent with higher levels of GSSG in the skin metabolism of the older group. This pattern implies that older individuals’ skin may undergo greater oxidative stress, as GSH is consumed to neutralize ROS, thereby increasing GSSG production. A larger absolute value of flux in the younger group also indicates a faster rate of GSSG reduction to GSH, suggesting that the skin microbiome of younger individuals may contribute to a stronger antioxidant recovery capacity and a more effective oxidative stress response, helping to maintain the balance of GSH/GSSG in the skin.

To identify key microbes and metabolites involved in the glutathione redox reaction, we applied shadow price analysis based on our constructed GEMs (**Table S5d**). Interestingly, *S. maltophilia*, which was more prevalent in the younger group, was predicted to utilize the glutathione cycle for antioxidant defense and maintenance of protein thiol homeostasis (**Fig. 4a,b**). It has been reported that *S. maltophilia* employs a variety of enzyme systems, including glutathione peroxidase, to combat oxidative stress from hydrogen peroxide, a type of ROS^75–77^. To determine whether the antioxidant defense capacity of *S. maltophilia* can mitigate cellular senescence in skin fibroblasts, skin primary fibroblasts were treated with H_2_O_2_ to simulate the oxidative stress which is typical of aged skin, followed by treatment with *S. maltophilia* culture supernatant (**Fig. 4c**). Results indicated that fibroblasts treated with *S. maltophilia* supernatant exhibited significantly increased levels of GSH and decreased levels of GSSG compared to the untreated control group. These effects are particularly evident in the H_2_O_2_-treated groups, where treatment with *S. maltophilia* supernatant leads to a markedly higher GSH/GSSG ratio (**Fig. 4d, Table S5e**). Given that senescent cells exhibit elevated β-galactosidase activity^78^, analysis using SA-β-gal staining shows that fibroblasts treated with *S. maltophilia* supernatant displayed reduced senescence markers (**Fig. 4e, Figure S4e**). This confirms the role of *S. maltophilia* in mitigating oxidative stress-induced cellular senescence.

**Fig. 4.**
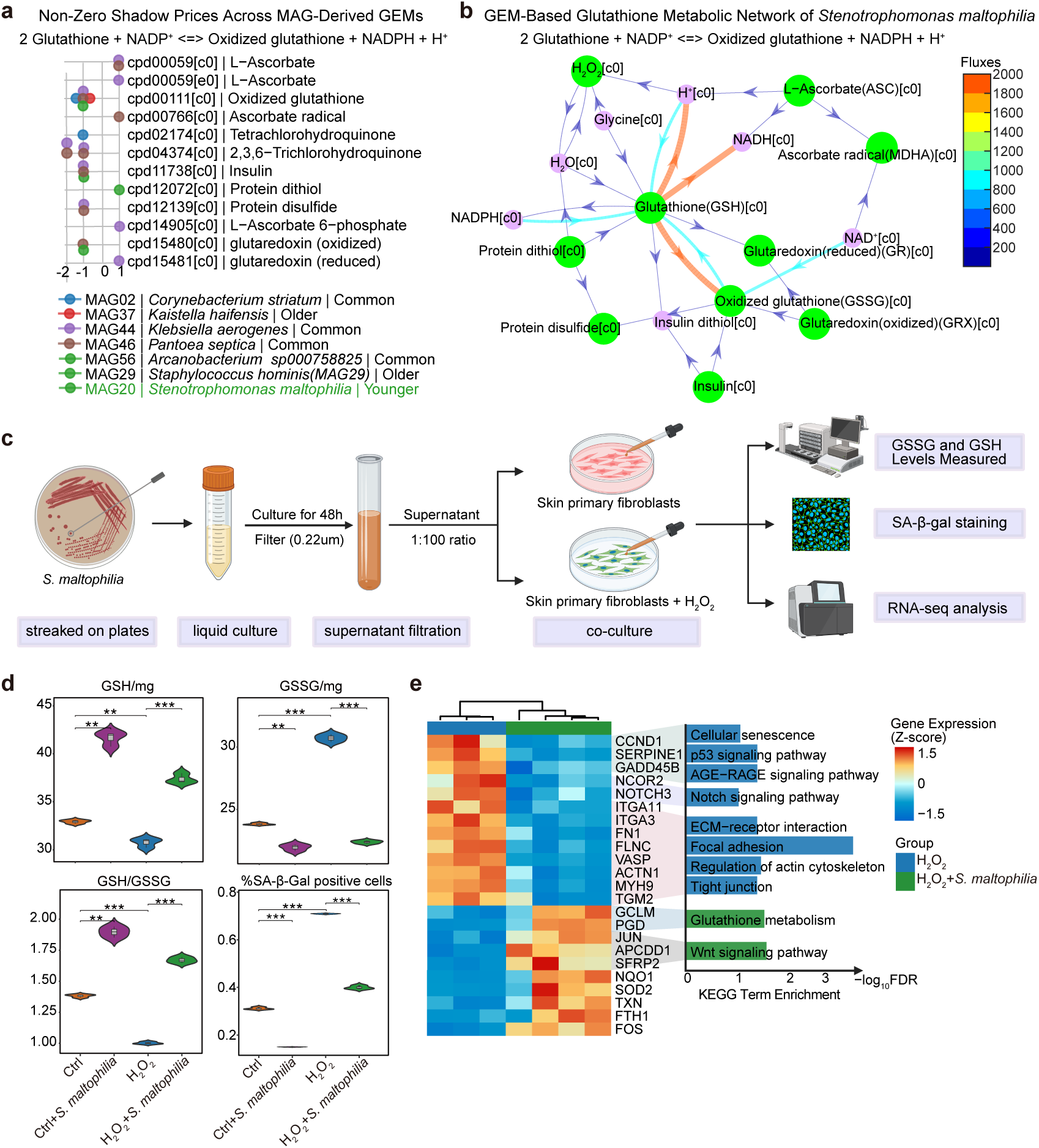
*S. maltophilia* in antioxidant defense. (a) Shadow price analysis from FBA in glutathione redox reactions across different MAGs. This plot illustrates the shadow prices from FBA related to glutathione oxidation-reduction reactions within the GEMs. The x-axis represents shadow price values, while the y-axis lists different metabolites. Positive values indicate that an increase in the metabolite concentration promotes the forward direction of the metabolic reaction, and vice versa. (b) Antioxidant defense network and flux analysis in *S. maltophilia* GEM: The Figure presents a network diagram of the glutathione (GSH)-dependent antioxidant defense mechanisms within the GEM of *S. maltophilia*. The diagram includes flux values depicting the dynamic metabolic interactions involving GSH, where it participates in reducing oxidized glutathione (GSSG) and combating oxidative stress via various enzymatic pathways, such as glutaredoxin. The fluxes are visualized in a color gradient representing their relative intensities, indicating the metabolic activity levels across different pathways. (c) Experimental workflow for investigating the effect of *S. maltophilia* supernatant on skin primary fibroblasts. *S. maltophilia* was first cultured on plates, followed by 48-hour liquid culture. The bacterial culture was filtered through a 0.22 µm filter, and the resulting supernatant was used to stimulate skin primary fibroblasts, both with and without H_2_O_2_ treatment, for 24 hours. The resulting samples were analyzed for GSH and GSSG levels, subjected to senescence-associated β-galactosidase (SA-β-Gal) staining for cellular senescence, and processed for RNA-seq analysis to assess gene expression changes. (d) Violin plots depicting the distribution of GSH, GSSG, GSH/GSSG ratio and the percentage of cells exhibiting SA-β-Gal activity in skin primary fibroblasts under different treatment conditions. Groups include control, control with *S. maltophilia*, H_2_O_2_ alone, and H_2_O_2_ with *S. maltophilia*. *P* values were calculated using T-test. Significance levels are indicated as follows: *, *P* < 0.05; **, *P* < 0.01; ***, *P* < 0.001. (e) Heatmap of differentially expressed genes between H_2_O_2_ alone, and H_2_O_2_ treatment combined with *S. maltophilia* culture supernatant. The left panel shows the Z-score normalized expression levels of the differentially expressed genes. The right panel displays the KEGG pathway enrichment analysis results for these genes (adjusted *P* < 0.1). Bar lengths represent the significance of enrichment, expressed as -log_10_(*p*.adjust), and are color-coded according to the treatment group.

To explore how *S. maltophilia* mitigates cellular senescence, we performed RNA-seq on H_2_O_2_-treated cells (**Table S5f**). Glutathione metabolism was significantly enriched in the *S. maltophilia* conditioned group (**Fig. 4e, Table S5h**), with upregulation of GCLM, enhancing GSH synthesis via glutamate-cysteine ligase activity^79^. PGD was also elevated, promoting NADPH supply for GSH regeneration^80^ (**Fig. 4e, Figure S4f**). These changes supported oxidative stress defense, maintained fibroblast viability, and delayed senescence, confirming that *S. maltophilia* alleviates oxidative stress by promoting GSH metabolism. Antioxidant genes *SOD2*, *NQO1*, *TXN*, and *FTH1* were significantly upregulated in the *S. maltophilia* group^81–84^ (**Fig. 4e**). Notably, TXN synergizes with GSH to maintain protein thiol redox balance, consistent with shadow price analysis indicating *S. maltophilia* supports thiol homeostasis (**Fig. 4a, Fig. S4c**). Meanwhile, the senescence-associated genes *SERPINE1, CCND1*, and *GADD45B* were downregulated^85^ (**Fig. 4e**). In contrast, pathways related to cell growth, regeneration, and tissue repair were enriched (**Fig. 4e, Fig. S4g**). Notably, high expression of *JUN*, *APCDD1*, and *SFRP2* activated Wnt signaling^86^, while downregulation of *NCOR2* and *NOTCH3*, and p53 signaling further promoted proliferation^85^ (**Fig. 4e, Fig. S4f,g**). Additionally, suppression of ECM remodeling and cytoskeletal pathways limited excessive matrix deposition, supporting tissue elasticity and flexibility^87^ (**Fig. 4e, Fig. S4g, Table S5i**).

In summary, our data support that the skin commensal *S. maltophilia* could effectively enhance antioxidant capacity and regenerative potential, thereby maintaining the viability of skin fibroblasts. In contrast, it might downregulate signaling pathways of cell cycle arrest, fibrosis, and accelerated senescence for skin cells. Although *S. maltophilia* is an opportunistic pathogen^39^, as previously mentioned, it is significantly associated with numerous youthful skin phenotypes (**Figure S3d**) and plays a role in both neutralizing oxidative agents and preserving protein function through thiol state maintenance, suggesting its potential as a target for anti-aging interventions.

### The potential anti-aging roles of *S. maltophilia* in Betaine, Lysolecithin, and Porphyrin metabolism

In addition to its role in alleviating oxidative stress-induced skin senescence, *S. maltophilia* demonstrates potential anti-aging properties through its involvement in other key metabolic pathways. These include the biosynthesis of betaine and lysolecithin, which are known for their skin-moisturizing, photoprotective, and anti-inflammatory properties, as well as its contributions to porphyrin metabolism, a process crucial for maintaining skin health and reducing oxidative damage (**Fig. 5, Figure S5**).

**Fig. 5.**
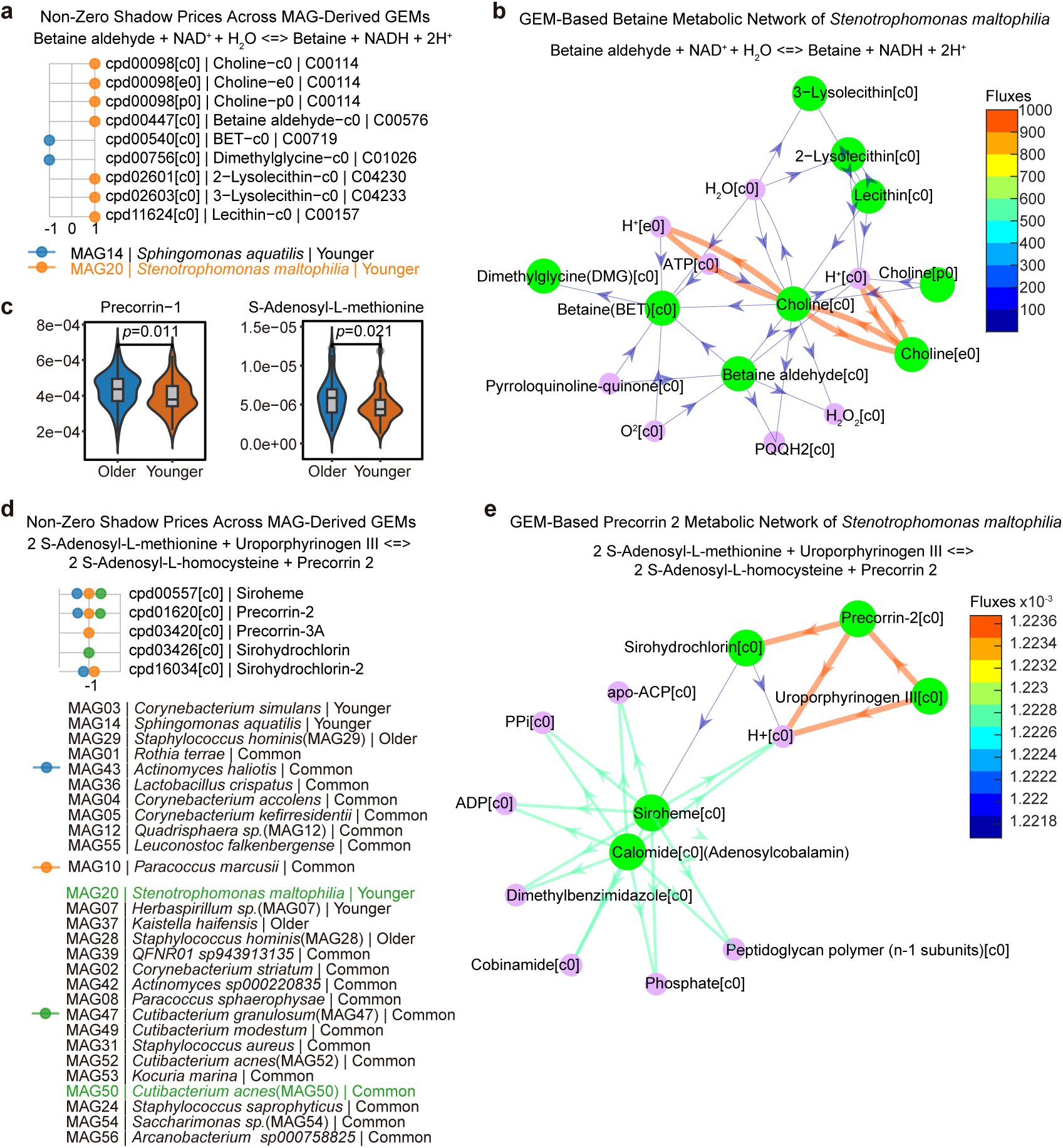
Vitamin B12 and Porphyrin biosynthesis pathways in skin microbiome and host co-metabolism. (a) FBA Shadow price analysis for betaine synthesis reactions across different MAGs. It highlights the biochemical value and metabolic impact of shifts in betaine-related compounds within microbial metabolism. The x-axis, y-axis, and color-coded dots follow a similar format to that in Fig. 4a. (b) Flux distribution diagram from the *S. maltophilia* GEM illustrating the betaine synthesis pathway. This diagram shows metabolic conversions, including the transformation of betaine aldehyde to betaine. The fluxes are visualized in a color gradient representing their relative intensities, indicating the metabolic activity levels across different pathways. (c) Violin plots illustrate significant differences in the concentrations of key porphyrin metabolism-related metabolites, such as precorrin-1 and S-Adenosyl-L-methionine, between older and younger groups. Notable differences are indicated, *P* < 0.05. (d) FBA Shadow price analysis for the conversion of Uroporphyrinogen III to Precorrin 2 in porphyrin metabolism. The x-axis, y-axis, and color-coded dots follow a similar format to that in Fig. 4a. (e) This Figure presents the GEM flux analysis of porphyrin metabolism pathways facilitated by *S. maltophilia*. It depicts how the conversion of Uroporphyrinogen III involves complex interactions leading to the synthesis of siroheme, among other compounds, with flux values shown in a color gradient to represent the intensity of metabolic activities.

Betaine and lysolecithin are known for their skin-moisturizing, photoprotective, and anti-inflammatory properties^88,89^. The biosynthetic pathways of betaine and lysolecithin have been reported in earlier studies^90,91^ (**Figure S5a**). Using GEMs, shadow price analysis of betaine and lysolecithin biosynthetic pathways identified *S. maltophilia* as a major microbial involved in their production in the skin (**Fig. 5a, Figure S5b, Tables S6a,b**). Furthermore, existing research suggests that lysolecithin may be a target or by-product of enzymatic activity in *S. maltophilia*^92,93^. Specifically, elevated shadow prices for precursors such as choline and betaine aldehyde in *S. maltophilia* suggest that increased availability of these substrates could enhance betaine production (**Fig. 5a**). Similarly, accumulation of intermediates in the acyl-CoA and fatty acid β-oxidation pathways may promote lysolecithin synthesis (**Figure S5b,c**). Exploring strategies to enhance the production of betaine and lysolecithin by *S. maltophilia* in the skin presents a potential direction for skin anti-aging.

Moreover, our previous analyses indicated that the younger group exhibits higher porphyrin optical density and porphyrin area percentage, along with a high enrichment of microbes involved in porphyrin metabolism (**Fig. 1b**, **Fig. 2e**). The porphyrin metabolic pathway, in which uroporphyrinogen III is converted into precorrin-2 and subsequently processed to form cobalamin (vitamin B12), has been well characterized in previous studies^94,95^ (**Figure S5d**). Through skin metabolomics analysis, we identified several metabolites involved in the porphyrin metabolic pathway (**Fig. 5c, Figure S5e**). Shadow price analysis performed on GEMs for the reaction converting Uroporphyrinogen III and Precorrin 2 revealed microbes associated with porphyrin metabolism (**Fig. 5d**). A key bacterium affecting this process is *C. acnes,* which has been reported to undergo transcriptional and metabolic changes in healthy individuals upon oral vitamin B12 supplementation^96^ (**Fig. 5d, Table S6c**). These changes include enhanced porphyrin production by *C. acnes*^96^. Elevated porphyrin accumulation could increase oxidative stress and pigment abnormalities in the skin^97^. However, adequate levels of Vitamin B12 facilitate rapid skin cell renewal and repair^98^, and provide antioxidant and photoprotective benefits^99^. Our data also underscore the significant role of *S. maltophilia* in the conversion of Uroporphyrinogen III to Precorrin 2 (**Fig. 5d, e**). Future research focusing on directing microbial utilization of body porphyrins towards Vitamin B12 synthesis, rather than permitting Vitamin B12 to be converted into intermediate porphyrins, could have profound implications for developing skin anti-aging strategies.

### Skin microbiota involvement in melanin metabolism during aging

The melanin synthesis pathway has been previously reported^100–102^. In this context, we elucidated a co-metabolic pathway of tyrosine metabolism shared between the skin microbiome and the host, which plays a crucial role in melanin synthesis within the skin (**Fig. 6a**). Our data showed that skin microorganisms in the younger group were significantly enriched in the tyrosine metabolism pathway (**Fig. 2d**). Utilizing established knowledge, the melanin biosynthesis process principally involves the conversion of Tyrosine to L-Dopa, followed by the oxidation of L-DOPA to Dopaquinone, which is then polymerized into melanin (**Fig. 6a**). Although no significant differences were observed in the levels of L-Tyrosine, DOPA, Dopaquinone, or Melanin within the skin metabolome, notable enrichment of L-Dopa, DA and 3-MT in the older group was detected (**Fig. 6b, Figure S6a**). Additionally, 3-MT exhibited a significant positive correlation with skin tone a* and b*, and a significant negative correlation with L* (**Fig. 3b**). These findings suggest that as the skin ages, DA and 3-MT may indirectly contribute to increased skin pigmentation. This highlights a potential role for these metabolites in the modulation of skin color and the aging process.

**Fig. 6.**
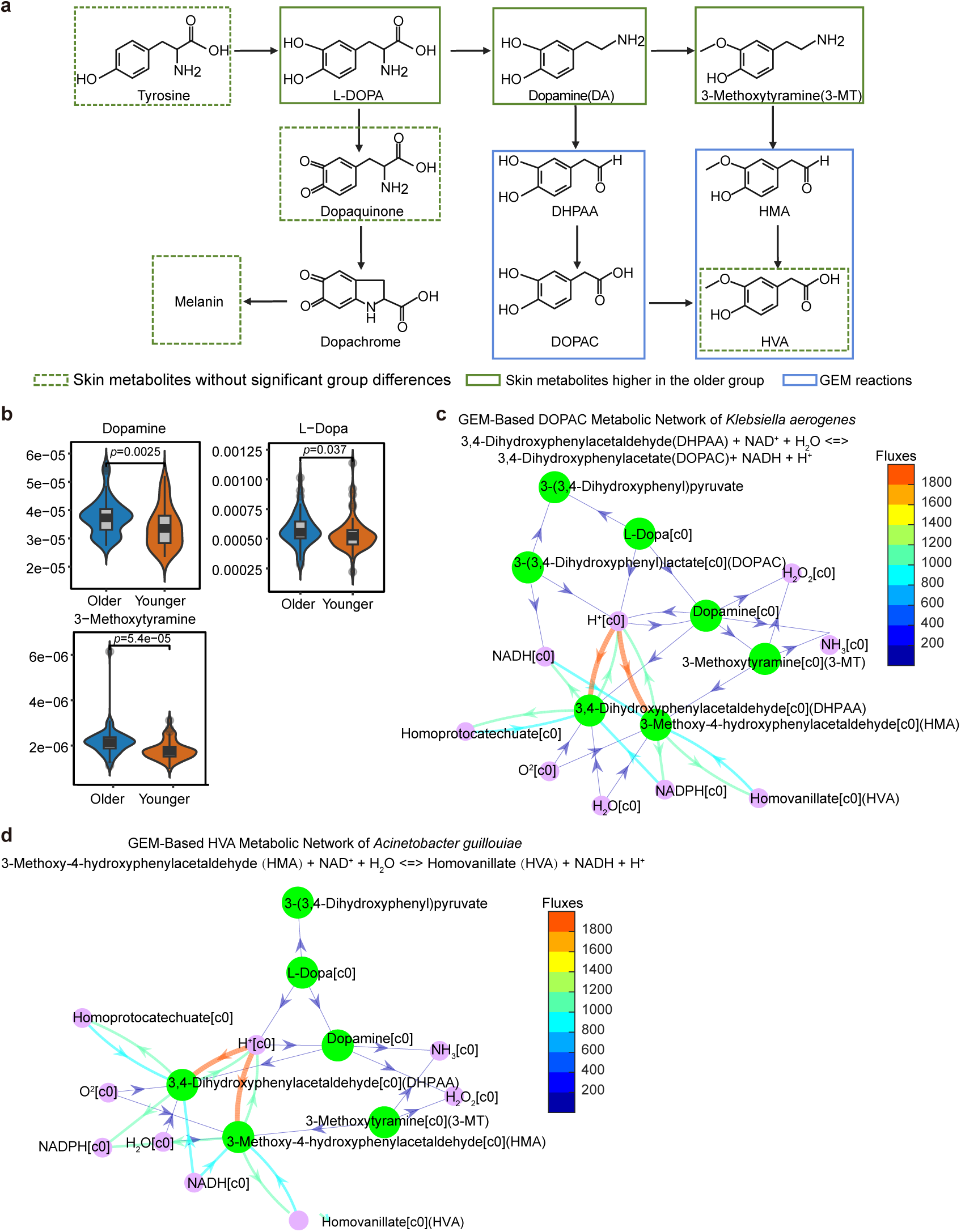
Tyrosine metabolism within the co-metabolic framework of skin microbiome and host. (a) Tyrosine metabolism in skin microbiome and host co-metabolism. This Figure illustrates the shared metabolic pathway of tyrosine between the microbiome and host, highlighting its role in producing key metabolites and melanin. Left Module (Tyrosine to L-3,4-dihydroxyphenylalanine (L-DOPA)): Tyrosine is hydroxylated to L-DOPA, a precursor for melanin and catecholamines, catalyzed by tyrosinase. L-DOPA is then oxidized to Dopaquinone, which cyclizes into Dopachrome and rearranges into Melanin. Right Module: L-DOPA is decarboxylated into dopamine (DA), which is metabolized into 3,4-Dihydroxyphenylacetaldehyde (DHPAA), 3,4-Dihydroxyphenylacetate (DOPAC), and Homovanillic acid (HVA). DA also forms 3-Methoxytyramine (3-MT), further processed into 3-Methoxy-4-hydroxyphenylacetaldehyde (HMA) and HVA. The diagram uses green dashed outlines to indicate skin metabolites that show no significant differences between groups, highlighting their consistent presence across different aging conditions. Green solid outlines denote metabolites with higher abundance in the older group, indicating changes that may be associated with aging. Blue outlines represent the reactions included in the GEM, emphasizing the biochemical interactions facilitated by the skin microbiome and host. (b) Violin plots depicting significant differences in key metabolites involved in tyrosine metabolism: Dopamine, L-Dopa, and 3-Methoxytyramine between older and younger groups in skin metabolomics. Notable differences are indicated, *P* < 0.05. (c) Tyrosine metabolism network and flux analysis in *K. aerogenes* GEM: This Figure illustrates the pathways involved in tyrosine metabolism within the GEM of *K. aerogenes*. The network shows the conversion of DHPAA to DOPAC and HMA to HVA. Flux values are indicated by a color gradient, representing the intensity of each reaction within the metabolic pathways. (d) Tyrosine metabolism network and flux analysis in *A. guillouiae* GEM: This Figure illustrates the pathways involved in tyrosine metabolism within the GEM of *A. guillouiae*.

GEM analysis demonstrated that the intermediate metabolic processes involving DA and 3-MT are influenced by the skin microbiota, particularly in the conversion pathways from DHPAA to DOPAC and from HMA to HVA (**Fig. 6a, Figure S6b,c, Tables S6d,e**). Shadow price analysis specifically highlighted the significant role of DA and 3-MT in regulating these microbial-involved metabolic pathways. A key microbe identified in this process is *Klebsiella aerogenes* (*K. aerogenes*). It has been reported to induce melanin production in fungi by providing melanin precursors^103^, thus acting as a microbial source of melanin substrates (**Fig. 6c, Figure S6b,c**). Interestingly, we also identified *A. guillouiae*, a microorganism enriched in the younger cohort, as a participant in this metabolic pathway, suggesting its potential capability in metabolizing melanin precursors (**Fig. 6d, Figure S3d, Figure S6b, c**). This suggests that exploiting microbial metabolism of DA and 3-MT to mitigate pigment deposition during skin aging presents a promising therapeutic avenue.

## Discussion

The relationship between skin aging and microbiome dynamics has been increasingly studied. Skin aging involves structural and physiological changes that correlate with shifts in microbial composition^22,24,32^. While prior studies focused on age-based population differences, our study targets individuals in their 40s, a critical watershed for skin aging, to investigate why some display significantly younger or older skin phenotypes. We aim to to uncover microbial regulatory mechanisms underlying this variation and identify strategies to slow aging through microbiome modulation.

Using a multi-omics approach integrating phenomics, metabolomics, metagenomics, and genome-scale metabolic modeling, we investigated skin aging within the same age group. Phenomics confirmed reliable stratification into younger and older phenotypes^12,13^. Notably, we found that younger skin phenotypes exhibited higher microbial diversity. This contrasts with earlier studies that reported lower diversity in youth, which we found was primarily due to the overwhelming abundance of *C. acnes* masking community richness^17,18,22,23,32^. Our findings suggest that enhanced microbial diversity supports skin homeostasis and a youthful appearance.

Our study reveals that the skin microbiome undergoes aging-related functional reprogramming, with older-associated communities favoring biosynthesis and genomic maintenance under nutrient-limited, stressed conditions^104^, while younger-associated microbiota support immune modulation and tissue renewal^51,105,106^. These microbial shifts are accompanied by metabolite differences: younger phenotypes are enriched in beneficial compounds with antioxidant and anti-inflammatory properties, such as methylcellulose^107^, alnustone^108^, and theobromine^109^, whereas pro-aging toxins like deoxynivalenol^62^ are more prevalent in older individuals. Predictive modeling revealed that combining microbiome and metabolomic data enhanced aging prediction, highlighting microbiome–metabolite interactions. This prompted GEM-based exploration of key microbial metabolites in skin aging.

*S. maltophilia*, typically known as an opportunistic pathogen^39^, has rarely been studied in the context of skin aging. Here, we provide the first evidence that, under commensal conditions, *S. maltophilia* contributes to redox homeostasis and skin health. GEM modeling and experimental validation showed that it enhances GSH synthesis and upregulates antioxidant genes such as *GCLM* and *SOD2*, improving fibroblast resistance to oxidative stress, a key driver of skin agin^72–74^. Beyond redox regulation, *S. maltophilia* may support barrier integrity and anti-inflammatory states by producing beneficial metabolites such as betaine and lysolecithin^88,89^. It also promotes vitamin B12 synthesis via porphyrin metabolism, potentially mitigating oxidative damage. These findings suggest a protective role for *S. maltophilia* in skin aging, warranting further investigation of its dynamic functions in the skin microenvironment. Our study also highlights the potential role of skin microbiota in melanin metabolism. *K. aerogenes* and *A. guillouiae*, enriched in younger individuals, may influence pigmentation via DA and 3-MT metabolism. While based on GEM modeling and correlation analysis, these findings warrant further functional validation in the context of skin aging.

In summary, our integrative multi-omics analysis provides critical insights into the intricate crosstalk between the skin microbiome and metabolites during the aging process, revealing how microbial-host interactions shape a youthful skin ecosystem. *S. maltophilia* supports antioxidant defense and hydration, while *A. guillouiae* may regulate melanin metabolism. Together, these findings highlight the therapeutic promise of targeting microbiome–metabolite crosstalk to delay skin aging, especially around the transitional age of the 40s.

Despite the strengths of our study, several limitations should be acknowledged. First, although we controlled for age range and geographic background, we did not collect detailed information on individual sun exposure, skincare practices, hormonal status, or lifestyle variables such as sleep and stress, all of which are known to influence both skin physiology and the microbiome. These unmeasured variables may introduce residual confounding. Future studies should incorporate these factors through comprehensive metadata collection and stratified sampling designs to further isolate the effects of the microbiome on skin aging.

## Data and code availability

The raw metagenomic sequencing data and RNA-seq data from this study are available in the NCBI BioProject database with accession number PRJNA1201908, and the raw metabolomics data have been deposited in the MetaboLights database with accession number MTBLS11994. Processed data matrices are provided in the Tables. The code used in this study is available on GitHub (https://github.com/guodanni1/Skin_Aging) to ensure reproducibility.

## Funding Declaration

This work was supported by Scientific Research Start-up Funds (QD2021005N), Shenzhen Science and Technology Program (WDZC20220819134430002), and Shanghai Jahwa United Co., Ltd.

## Author contributions

Xiao Liu and H.J. conceived the idea; Xiao Liu and Haidong Jia supervised the work; H.J., Xiao Liu, J.C., Y.C. designed the experiments; Danni Guo wrote the manuscript; J.C. and W.L. collected the samples with the help of Y.C.; Jingmin Cheng performed the majority of the experiments with the help of W.M., H.Y.; Danni Guo analyzed the data with the help of J.C.; Yahong Wu conducted the functional experiments; Xiao Liu, L.H., and L.M. reviewed and edited the manuscript.

## Declaration of interests

Haidong Jia and Yuanyuan Chen receive salaries from Shanghai Jahwa United Co., Ltd., and the other authors declare no competing interests.

## Materials and methods

### Sample collection

We initially recruited 202 healthy Chinese women aged 39 to 41 from the general population in Shanghai in August 2022. This specific age range was chosen to capture a well-defined transitional window, as there is growing evidence that the 40s represent a critical inflection point in skin aging^27–29^, during which biological and structural changes become more pronounced. To further minimize confounding factors that could influence the skin microbiome, all participants were recruited from the same geographic region and were expected to share similar genetic backgrounds, dietary habits, and lifestyle patterns. Individuals with a history of skin diseases, recent anti-aging treatments, or antibiotic use within the past month were excluded. To maximize microbial skin load, all participants were instructed to avoid facial cleansing or the use of any skincare or cosmetic products within 12 hours prior to sampling. Specifically, subjects were asked to perform their final facial wash using only tap water before 8:00 p.m. on the evening prior to sampling and to refrain from washing their face the following morning.

Ethics approval for this study was obtained from the Ethics Committee of Tsinghua Shenzhen International Graduate School (Approval No. [2022] 97). Written informed consent was obtained from all participants before skin microbiome sampling by institutional guidelines.

### Skin phenotype assessment

We evaluated skin characteristics in 202 healthy women, focusing on ecological niche indicators and appearance. Traits measured included transepidermal water loss (TEWL), stratum corneum water content (WCSC), sebum production, surface pH, elasticity, and porphyrins (area percentage, average area, and optical density), as well as wrinkles and skin tone (L*, a*, b*). TEWL, reflecting skin barrier function, was gauged using a Vapometer^®^ (Delfin Technologies Ltd). WCSC was determined via a Corneometer^®^ CM 825 (EnviroDerm Services, UK), and sebum output measured with a Sebumeter^®^ SM815 (Courage & Khazaka electronic GmbH), noted in μg/cm^2^. Skin pH was assessed using a Skin-pH-Meter PH 900 (Courage & Khazaka electronic GmbH). Elasticity, indicated by the R2 parameter (ratio of retraction to deformation), was analyzed with a Cutometer^®^ dual MPA 580 (Courage & Khazaka electronic GmbH). Skin tone, porphyrins metrics, and porphyrin distribution were analyzed using ImageJ, based on VISIA-CR images (Canfield Scientific Inc). Wrinkle assessment was conducted by dermatologists. They examined various types including forehead lines, crow’s feet, glabellar, under-eye wrinkles, and nasolabial folds, with a composite score determining overall wrinkle severity.

### Grouping criteria and sample collection procedures

To stratify participants based on apparent skin aging, we applied an artificial intelligence-based facial analysis platform (Megvii Face++)^30^, which estimated each participant’s perceived age. Based on this assessment, 100 women whose AI-estimated ages were significantly lower than their chronological age were classified as the younger group, and 102 whose AI-estimated ages were significantly higher were classified as the older group.

In addition to AI-based grouping, we used R2 (gross elasticity) as a complementary criterion to identify individuals with distinct biological aging profiles, as it reflects dermal properties that are not readily captured by AI-estimated age based on facial images (Megvii Face++^30^). For downstream analysis, we selected a subset of 103 participants, including the 52 individuals from the younger group with the highest R2 scores and the 51 from the older group with the lowest R2 scores.

To maximize skin microbial yield, 103 human subjects were instructed to wash their faces only with tap water and avoid applying any skincare or makeup products prior to sampling. The left cheek was sampled for metagenomic sequencing, and the right cheek for metabolomic analysis. Sterile medical cotton swabs pre-moistened with 0.15 M NaCl solution were used to swab each area 25 times, utilizing a systematic vertical and horizontal motion for thorough sampling. The swab heads were then detached and stored in sterilized 1.5 mL centrifuge tubes at −80°C. A total of 103 facial microbial samples were collected.

### DNA extraction and metagenomic sequencing

The experimental workflow encompassed several key stages: DNA extraction, DNA quality assessment, library preparation, library quantification, and sequencing. DNA extraction was carried out using the Novizan FastPure^®^ Host Removal and Microbiome DNA Isolation Kit (item No.: DC501), adhering strictly to the manufacturer’s protocol. This kit is specifically designed to efficiently lyse host cells and deplete host genomic DNA, while preserving microbial DNA from skin swabs. DNA concentration was determined using the Qubit fluorometer. Subsequent library preparation was conducted utilizing the Novizan VAHTS^®^ Universal Plus DNA Library Prep Kit for MGI (item No.: NDM617). Post-preparation, libraries were initially quantified with the Qubit fluorometer and then diluted to a concentration of approximately 5 ng/µl to optimize sequencing conditions. Each step underwent rigorous quality control to ensure reliability, and sequencing was performed on an MGI platform using the PE150 strategy to a depth of approximately 12 Gbp each sample.

### Preparation of metabolomics samples and LC-MS measurement

All chemicals and solvents utilized were of analytical or HPLC grade. Water, methanol, acetonitrile, and formic acid were acquired from Thermo Fisher Scientific (Waltham, MA, USA). L-2-chlorophenylalanine was sourced from Shanghai Hengchuang Bio-technology Co., Ltd. (Shanghai, China), and chloroform was obtained from Titan Chemical Reagent Co., Ltd. (Shanghai, China). Each sample was treated with a total of 2 mL of pre-chilled methanol-water mixture (4:1, v/v), which was transferred in two portions to glass vials. The samples were then subjected to ultrasonication for 20 minutes in an ice-water bath. After extraction, samples were allowed to stand at −40°C for 30 minutes before being centrifuged at 13,000 rpm for 10 minutes at 4°C. 1mL of the supernatant was then transferred to an LC-MS sample vial and dried. For reconstitution, 300 µL of methanol-water mixture (1:4, v/v) containing 2 µg/mL of L-2-chlorophenylalanine was added to each sample. Samples were vortexed for 30 seconds, ultrasonicated for 3 minutes in an ice-water bath, and subsequently rested at −40°C for 2 hours. Post-resting, samples were centrifuged at 4°C (13,000 rpm) for 10 minutes. A 150 µL aliquot of the supernatant was drawn with a crystal syringe, filtered through a 0.22 µm organic phase syringe filter, and transferred into LC vials. The vials were stored at - 80°C until LC-MS analysis. Quality control (QC) samples were prepared by pooling equal volumes of the extracts from all samples. All reagents used in the extraction process were pre-chilled to −20°C prior to use.

Metabolomic profiling was performed using an ACQUITY UPLC I-Class system coupled with a VION IMS QTOF Mass Spectrometer (Waters Corporation, Milford, USA). The analyses were conducted in both ESI positive and ESI negative ion modes to enhance the detection spectrum. Chromatographic separation was achieved on an ACQUITY UPLC HSS T3 column (100 mm × 2.1 mm, 1.8 µm). The mobile phases consisted of water (A) and a mixture of acetonitrile/methanol (2:3 v/v) (B), both containing 0.1% formic acid. A linear gradient was employed as follows: 0 min, 5% B; 2min, 5% B; 4min, 30% B; 8min, 50% B; 10min, 80%B; 14 min, 100% B; 15 min, 100% B; 15.1 min, 5% B and 16min, 5%B. The flow rate was maintained at 0.35 mL/min, and the column temperature was set at 45°C. Sample injection volume was 2 µL, and samples were kept at 4°C during the analysis.

Mass spectrometry was performed in full scan mode, covering a mass-to-charge (m/z) range of 100 to 1200. Data acquisition was supplemented by MSE mode, where two independent scans with varying collision energies were alternated: a low-energy scan at 4 eV and a high-energy scan with a ramp from 20 to 45 eV for ion fragmentation. The spectrometer was operated with the following parameters: resolution (full scan): 70,000, resolution (HCD MS/MS scans): 17,500, spray voltage: 3.8 kV for positive mode and −3.0 kV for negative mode, sheath gas flow rate: 35 Arb, aux gas flow rate: 8 Arb, capillary temperature: 320°C, argon (99.999%) as the collision gas, a scan time of 0.2 seconds, an interscan delay of 0.02 seconds, a capillary voltage of 2.5 kV, a cone voltage of 40 V, a source temperature of 115°C, a desolvation gas temperature of 450°C, and a desolvation gas flow of 900 L/h. Quality control samples were injected at regular intervals, after every 10 samples, to assess the repeatability and stability of the analytical conditions throughout the run.

### LC-MS data processing and metabolomics analysis

LC-MS raw data were processed using Progenesis QI v2.3 software (Nonlinear Dynamics, Newcastle, UK). The analysis involved baseline filtering, peak identification, integration, retention time correction, peak alignment, and normalization. Normalization of metabolite intensities was conducted using the total ion current (TIC) method to correct for variations in signal intensities across samples. Peak detection and alignment were performed using the default apex peak picking algorithm in Progenesis QI. The analysis settings included a precursor ion mass tolerance of 5 ppm, a product ion mass tolerance of 10 ppm, and a minimum product ion threshold of 5%. Compound identification was performed based on accurate mass-to-charge ratio (m/z), secondary fragmentation patterns, and isotopic distributions. The identification databases utilized included the Human Metabolome Database (HMDB)^110^, LipidMaps v2.3^111^, METLIN^112^, EMDB^113^, PMDB^114^, and a proprietary database developed by Shanghai OE Biotech Co., Ltd., facilitating comprehensive qualitative analysis. Data refinement involved the exclusion of any peaks that exhibited a missing value (ion intensity = 0) in more than 50% of the samples within any group. To ensure analytical reproducibility, features with a relative standard deviation (RSD) >30% in pooled QC samples were removed from subsequent analyses. Zero values were imputed using half of the minimum detected value to maintain data integrity. Compounds were screened based on their qualitative identification scores, and those scoring below 36 out of a possible 60 were excluded from further analysis to ensure data accuracy. Finally, a consolidated data matrix was created by integrating the positive and negative ionization data results. Additionally, data were analyzed using MetOrigin to identify enriched pathways.

### Quality control and contaminant removal of metagenomic reads

Raw paired-end metagenomic reads were subjected to a comprehensive quality control (QC) pipeline using BBTools (v39.01). Initial reformatting was performed using ReformatReads to standardize sequence headers, convert all IUPAC ambiguity codes to “N” (iupacToN=t), enforce uppercase sequences (touppercase=t), and ensure consistent Phred+33 quality encoding (qout=33). Read pairing integrity was verified with verifypaired=t, and the output reads were saved as compressed FASTQ files.

To eliminate PCR or optical duplicates, we applied Clumpify with duplicate removal enabled (dedupe=t), allowing a maximum of two mismatches per duplicate pair (dupesubs=2). This step effectively collapsed redundant read clusters, yielding a deduplicated dataset for downstream analysis.

Quality trimming and adapter removal were performed using BBDuk. Adapter sequences were trimmed from the 3′ ends of reads (ktrim=r) using a k-mer size of 27 (k=27) with a minimum match of 8 bp (mink=8) and allowing one mismatch (hdist=1). Both ends of reads were trimmed for low-quality bases (qtrim=rl) using a Phred threshold of 10 (trimq=10), and reads shorter than 51 bp after trimming were discarded (minlength=51). Overlapping paired-end reads were subjected to error correction (ecco=t), and multi-threaded parallelization was enabled (pigz=t, unpigz=t) with memory preallocation (prealloc=t) for performance optimization.

Following quality filtering, host and contaminant sequences were removed using BBSplit, aligning reads against a custom merged reference containing host (e.g., human), PhiX, and other potential contaminant genomes. The alignment was conducted in local mode (local=t) with a relaxed k-mer size of 13 (k=13) and maximum allowed indels of 20 (maxindel=20). Reads were retained if they exhibited at least one matching k-mer (minhits=1) and a minimum alignment score ratio of 0.65 (minratio=0.65). Ambiguously mapped reads were resolved by selecting the best hit (ambiguous=best). Clean reads not mapping to any contaminant reference were output as final QC-passed datasets. Only high-quality, de-duplicated, non-host reads were retained for downstream taxonomic and functional profiling.

### Metagenomic data assembly

Assembly of metagenomic sequences were conducted using the Metagenome-atlas v2.16.3 pipeline^115^. The process involved the ‘atlas run all’ command for streamlined execution. Initially, adapter sequences were trimmed, and human genomic contaminations were removed utilizing BBMap suite v39^116^. Subsequently, reads underwent error correction and were merged prior to assembly with MEGAHIT v1.2.9^117^. For binning, MetaBAT v2.15^118^ and MaxBin v2.2^119^ were employed, and the integrated results were processed using DAS Tool v1.1^120^. The resultant MAGs, which achieved a minimum of 50% completeness and less than 10% contamination as assessed by CheckM v2^121^, were clustered based on 97.5% average nucleotide identity. 50 species-representative MAGs and 6 strain-level MAGs were then taxonomically annotated using GTDB-tk v2.1 against Genome Taxonomy Database release 214.0^122^. The resulting phylogenetic tree was visualized using Interactive Tree Of Life (iTOL) v6^123^.

### Taxonomic identification and functional profiling of microbial communities

High-quality reads from each sample were mapped to a reference set of 2,419 species-level microbial Operational Taxonomic Units (mOTUs) for taxonomic identification^124^. Functional profiling was conducted using HUMAnN v3.6^125^ with default parameters for all modules. Differentially enriched KEGG modules were determined based on their reporter scores. These scores were calculated from the Z-scores of individual KEGG Orthologs (KOs) by the “ReporterScore” R package v0.1.4^126,127^. A detection threshold was set at an absolute reporter score of 1.96 to identify modules showing significant differences in abundance.

### Construction of Random Forest models for skin aging prediction

To evaluate the predictive performance of metabolomics, microbiome, and combined omics data in skin aging, we constructed Random Forest models. All models were implemented using the randomForest package v4.7-1.2^128^ and reproducibility was ensured with a fixed random seed (set.seed(123)). The datasets were initially split into training and validation sets using a 7:3 ratio via stratified sampling with the createDataPartition() function, ensuring consistency of sample assignments across all three groups (metabolites, microbials, and combined omics) to maintain comparability across models. The original, unfiltered features consisted of 247 microbial species (selected based on frequency > 30% and relative abundance > 0.01), 3,033 core metabolites (selected based on frequency > 30% and relative abundance > 0.001%). Feature importance was evaluated using the MeanDecreaseAccuracy metric derived exclusively from the training dataset via the importance() function. Significant features for each data type metabolites, microbials, and combined microbials + metabolites were independently selected using this metric. For each group, features with a MeanDecreaseAccuracy > 1 were retained for downstream analysis. A new Random Forest model was trained using these significant features, with mtry set to the square root of the feature count and ntree fixed at 200. Model performance on validation datasets was assessed using sensitivity, specificity, precision, recall, F1 score, and area under the ROC curve (AUC). The caret package v7.0-1^129^ was used for confusion matrix computation and performance evaluation, while the pROC package v1.18.5^130^ was employed for ROC curve generation and AUC calculation. Data visualization, including ROC curves, was conducted using ggplot2 v3.4.4^131^, ggpubr v0.6.0, and ggprism v1.0.5.

### Genome-scale metabolic model (GEM) construction

In this study, GEMs were constructed based on metagenomic data from 103 samples, which produced 56 MAGs using the Metagenome-Atlas v2.16.3^115^ pipeline as mentioned above. The genome sequences of these MAGs were stored in .fna files for our GEM construction. We utilized gapseq v1.2^132^, a tool designed for the efficient and robust construction of metabolic pathways and the identification of transport proteins through homology-based methods. The ‘gapseq doall’ process was instrumental in our approach, encompassing the identification of metabolic pathways (‘find’ function) and transport proteins (‘find-transport’ function), followed by the generation of preliminary draft models. These models were further refined by a thorough gap-filling process (‘fill’ function) to address metabolic inconsistencies. To generate individual sample-specific GEMs, we employed the “mergeTwoModels” function alongside our custom-developed function, “processDuplicateReactions”. The methodology involved weighting the flux bounds of reactions in each MAG GEM based on the relative abundance of the corresponding MAG in each sample. This process ensured that the flux bounds of the same reactions across different MAGs within a sample were aggregated based on their weighted values to establish the flux bounds in the sample’s GEM. Subsequently, these sample-specific GEMs were merged according to group classifications (older or younger), forming two distinct group-specific GEMs.

### Metabolic network analysis

For analyzing and manipulating the reconstructed metabolic models, we employed the COBRA Toolbox (2024 release)^133^, a MATLAB toolbox used for constraint-based modeling of metabolic networks. We performed linear and integer programming simulations using the Gurobi Optimizer (version 1003)^134^. Flux Balance Analysis (FBA) of each reaction was conducted using the ‘optimizeCbModel’ function of the COBRA Toolbox, which optimizes metabolic networks to elucidate metabolic capabilities. We also assessed the impact of metabolites and microbes on reactions by computing shadow prices using ‘FBAsolution.f’ and ‘FBAsolution.y’ functions from the toolbox. For visualizing metabolic networks, the ‘createMetIntrcNetwork’ function from the COBRA Toolbox was used to generate interactive network graphs, highlighting the interconnectivity and flux distributions within our metabolic models, thus providing a detailed view of the metabolic interactions.

### Statistical analysis and data visualization

Differences in skin phenotypes, microbial species, and metabolites between older and younger groups were evaluated using Wilcoxon rank-sum tests to identify statistically significant variations. The *p-*values derived from these tests were adjusted for multiple comparisons employing the False Discovery Rate (FDR) correction method. Microbial differential abundance analysis was specifically performed on taxonomic profiles using Wilcoxon rank-sum test, and the resulting p-values were adjusted for multiple comparisons using the Benjamini-Hochberg FDR method (adjusted *P* < 0.05 considered significant), and biological relevance (absolute log_2_fold change > 1) was also required to define differential abundance. This adjustment was conducted using the “stats” package in R v4.1.1. Additionally, correlations were assessed using Spearman’s rank correlation coefficient, implemented through the “psych” package v2.3.3^135^ in R, to explore relationships between variables across different groups. Python v3.8.8 and the Pandas library v1.2.4^136^ were utilized for compiling shadow price analysis results. The R packages ggplot2 v3.4.4^131^ and ggrepel v0.9.3^137^ facilitating the data visualization.

### Bacterial growth conditions and supernatant preparation

The *Stenotrophomonas maltophilia* strain used in this study was obtained from the BeNa Culture Collection (BNCC) under preservation number BNCC185982. The *S. maltophilia* strain was streaked onto Brain-Heart Infusion (BHI) agar plates and incubated overnight at 37°C. The following day, individual colonies were inoculated into liquid BHI medium and cultured overnight at 37°C with shaking at 220 RPM. Subsequently, cultures of the activated strain were transferred into fresh BHI medium. After 24 hours of incubation, when cultures reached an OD_600_ of 0.8-1.0, bacterial cells were pelleted by centrifugation, and the culture supernatants were filtered twice through 0.22 μm Spin-X centrifuge tube filters (Corning). The supernatants were then stored at −80°C for subsequent experiments. Notably, supernatants were also collected at 12 hours and 72 hours post-inoculation, respectively, to achieve the desired OD_600_ values.

### Cell culture

Primary human skin fibroblast cultures were derived from punch biopsies collected from healthy male donors undergoing circumcision surgery, with informed consent obtained according to ethical guidelines. Briefly, skin specimens were immersed in sterile normal saline and washed three times with phosphate-buffered saline (PBS). The specimens were then cut into 1 mm × 1 mm sections and placed in a 6-cm Petri dish, with the epidermis facing upward and the dermis downward. Dulbecco’s Modified Eagle Medium (DMEM) supplemented with 10% (v/v) fetal bovine serum (FBS) and 1% (v/v) penicillin-streptomycin (all from Gibco, USA) was added to each dish. The culture medium was replaced every 2-3 days. After 7-14 days, dense fibroblast outgrowths were observed and subsequently passaged using 0.25% trypsin-EDTA. All experiments utilized primary fibroblasts within five passages. Cultures were maintained in a standard incubator at 37°C with 5% CO_2_.

### Stimulation of fibroblasts with bacterial supernatants

Primary human skin fibroblasts were plated in 6 cm Petri dishes and grown until reaching approximately 80% confluence. Filtered supernatants from *S. maltophilia* cultures were diluted in DMEM at a ratio of 1:100 and used to stimulate fibroblasts for 24 hours. Cells treated with BHI alone served as the control group.

### Quantification of GSH and GSSG levels in fibroblasts

GSH and GSSG levels in human skin fibroblasts were quantified using the S0053 Glutathione Assay Kit (Beyotime, China) following the manufacturer’s protocol. Cell samples were lysed to release intracellular glutathione. Total glutathione content, comprising both reduced (GSH) and oxidized (GSSG) forms, was measured by reducing GSSG to GSH using glutathione reductase. The resulting GSH reacted with 5,5’-dithiobis-(2-nitrobenzoic acid) (DTNB), forming the yellow compound 5-thio-2-nitrobenzoic acid (TNB). The absorbance of TNB was measured at 412 nm (A412) to quantify total glutathione. To specifically measure GSSG levels, a GSH masking reagent provided in the kit was used to selectively remove GSH from the samples before repeating the reduction and detection process. The concentration of reduced GSH was calculated by subtracting the GSSG concentration from the total glutathione content.

### β-Galactosidase staining for cellular senescence detection

Cellular senescence was assessed using the β-Galactosidase Staining Kit (Cat # KTA3030) following the manufacturer’s protocol. Adherent cells were cultured in 24-well plates and subjected to the appropriate treatments. After treatment, the culture medium was aspirated, and the cells were washed twice with 1× PBS. Next, 1 mL of 1× fixative solution was added to each well, and the cells were fixed at room temperature for 15 minutes. Following fixation, the fixative solution was removed, and the cells were washed three times with 1× PBS. Subsequently, 1 mL of staining working solution was added to each well, and the plates were incubated overnight in a CO_2_-free incubator at 37°C until the cells developed blue staining. To prevent cell drying during incubation, the plates were sealed with parafilm, and the staining duration was adjusted to achieve optimal staining, the cells were observed under a light microscope to evaluate β-galactosidase activity.

### RNA Sequencing

Total RNA was extracted from primary fibroblasts using the RNA Quick Extraction Kit (Magen, China) according to the manufacturer’s protocol. To remove DNA contamination, total RNA samples were treated with DNase I (NEB), and the reaction was terminated by adding 0.5M EDTA, followed by incubation at 75°C for 10 minutes and cooling on ice. The samples were then purified using RNA Clean beads (Vazyme). Enrichment and purification of mRNA were performed using mRNA Capture Beads (Vazyme). The enriched mRNA was incubated with 5× HiScript II Buffer (Vazyme) and N6 primer (0.1 µg/µL) at 94°C for 10 minutes. The fragmented RNA was combined with dNTP mix (10 mM, Vazyme), RNase Inhibitor (40 U/µL, Vazyme), and HiScript II Reverse Transcriptase (200 U/µL, Vazyme) for reverse transcription. The first-strand reaction products were added to a mix containing 5×Second Strand Buffer (Invitrogen), dNTP mix (10 mM, Vazyme), RNase H (5 U/µL, Yeasen), DNA Polymerase I (5 U/µL, Takara), and nuclease-free water (Invitrogen). The reaction was carried out at 16°C for 2 hours, and the samples were purified using DNA Clean Beads (Vazyme). End repair and A-tailing were performed using T4 DNA Polymerase, Klenow DNA Polymerase, and T4 Polynucleotide Kinase (Vazyme). dA tails were added using Taq DNA Polymerase (Vazyme). Library amplification was performed using AD153 primers and 2×KAPA HiFi HotStart ReadyMix (Vazyme). The final libraries were purified, quantified, and pooled for sequencing. Paired-end sequencing (PE150) was conducted on the MGI T7 platform. The integrity of RNA was assessed using the RNA Nano 6000 Assay Kit on the Agilent Bioanalyzer 2100 system (Agilent Technologies, CA, USA). The quality of libraries was also verified using the Agilent Bioanalyzer 2100 system.

### Processing and analysis of RNA-seq data

Raw reads obtained from the high-throughput sequencing platform were processed to generate an expression matrix through the following steps. Quality control of the raw data was conducted using FastQC v0.12.1^138^ to assess sequencing quality. Low-quality reads and adapter sequences were removed using Trim Galore v0.6.10^139^, and non-coding RNA sequences were filtered out using SortMeRNA v4.3.6^140^. The cleaned reads were aligned to the reference genome using STAR v2.7.10b^141^, and gene-level read counts were quantified by mapping the alignment results to gene annotations with FeatureCounts v2.0.6^142^. The final expression matrix served as the input for subsequent differential expression and downstream analyses. Differential expression analysis was performed using the DESeq2 R package v1.32.0^143^, identifying genes with an adjusted *P* < 0.05 and absolute Log_2_FoldChange > 0.5 as differentially expressed. Gene Ontology (GO) and KEGG enrichment analyses were conducted using the clusterProfiler R package v4.0.5^144^, with KEGG terms considered significantly enriched at an adjusted *P* < 0.1.

