## Supplementary Figures 1-6 for "Multi-omics characterization of the skin microbiota reveals the anti-aging roles of *Stenotrophomonas maltophilia*"

### Supplementary figures and figure legends

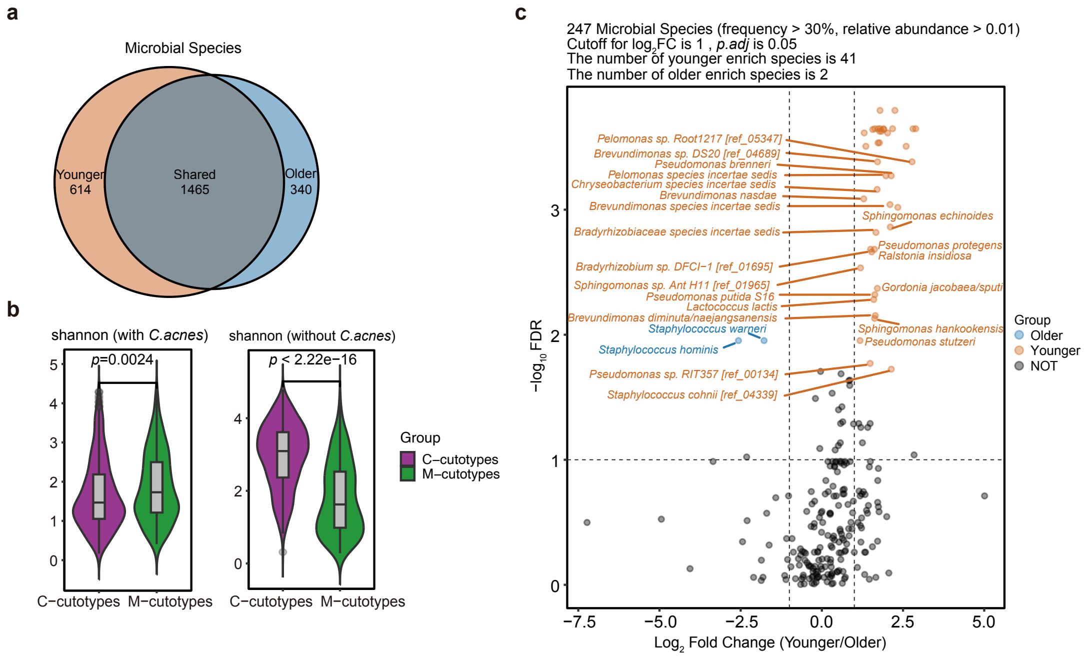


**Figure S1. Metagenomic analysis reveals skin aging-related microbial dynamics.**

(a) Venn diagram displays the distribution of microbial species identified in metagenomic analysis.

(b) Violin plots show microbial diversity with and without the influence of C. acnes.

(c) Volcano plot illustrates the differential enrichment of 43 microbial species between groups. Volcano plot of microbial species enriched in younger and older skin microbiomes based on differential abundance analysis using the Wilcoxon rank-sum test. The analysis included 247 microbial species with a frequency >30% and relative abundance >0.01. The analysis used an absolute $Log_2FoldChange > 1$ and adjusted $P < 0.05$.

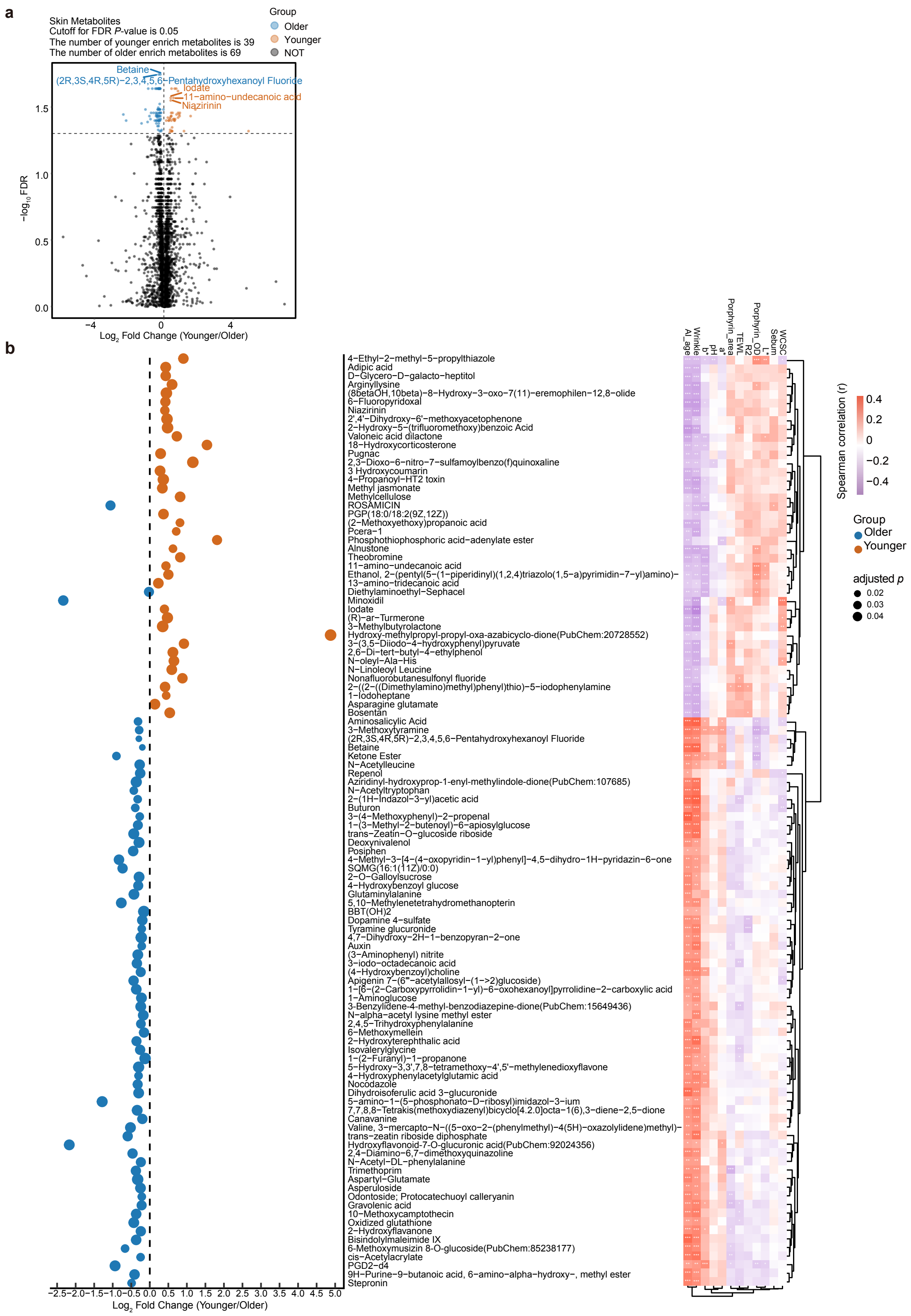


**Figure S2. Metabolomic analysis reveals skin aging-related metabolites.**

(a) Volcano plot illustrates the differential enrichment of 108 skin metabolites between groups.

(b) Left: Bubble chart displays all skin metabolites with significant differences between groups (adjusted $P < 0.05$). Right: Heatmap shows the Spearman correlation analysis between all differential skin metabolites and skin phenotypes. Significance levels are indicated as follows: *, adjusted $P < 0.1$; **, adjusted $P < 0.05$; ***, adjusted $P < 0.01$.

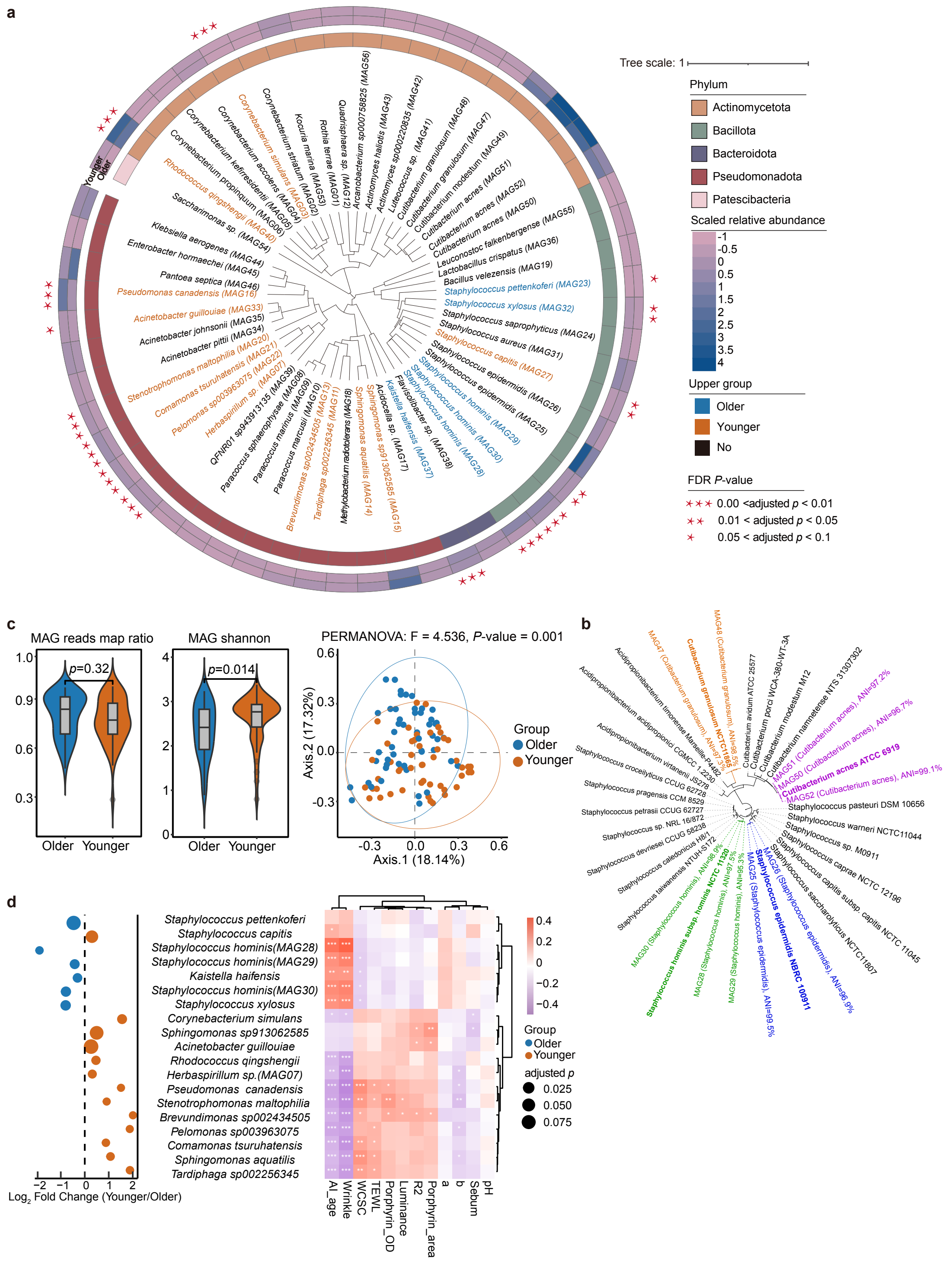


**Figure S3. MAGs analysis of age-related differences in skin microbiota.**

(a) Phylogenetic tree with heatmap: This visualization combines a phylogenetic tree with a heatmap representing 56 MAGs derived from 103 samples. Microbial names in blue indicate MAGs with higher abundance in the older group, while those in orange represent higher abundance in the younger group. From the center outward, the first circle indicates the phylum, the second circle shows relative abundance in the older group, the third circle shows relative abundance in the younger group, and the outermost stars indicate significance levels: *, adjusted $P < 0.1$; **, adjusted $P < 0.05$; ***, adjusted $P < 0.01$.

(b) Phylogenetic tree of strain variation: This tree displays different strains of the same species and their positions within the annotated GTDB database. The Average Nucleotide Identity (ANI) values indicate the genetic similarity of each strain to its nearest annotated counterpart.

(c) Left: Violin plots illustrate the ratio of reads mapped to MAGs versus the total clean reads, suggesting that the MAGs can represent a significant portion of the microbial community in both age groups without significant differences. Middle: Violin plots display microbial diversity calculated using the relative abundance of MAGs. Right: PCA plots highlight significant differences in the compositional structure of the MAGs between the groups, underlining the distinct microbial community structures associated with different age groups.

(d) Left panel: Bubble chart displays all MAGs with significant differences between groups (adjusted $P < 0.05$). Right panel: Heatmap shows the Spearman correlation analysis between all differential MAGs and skin phenotypes. Significance levels are indicated as follows: *, adjusted $P < 0.1$; **, adjusted $P < 0.05$; ***, adjusted $P < 0.01$.

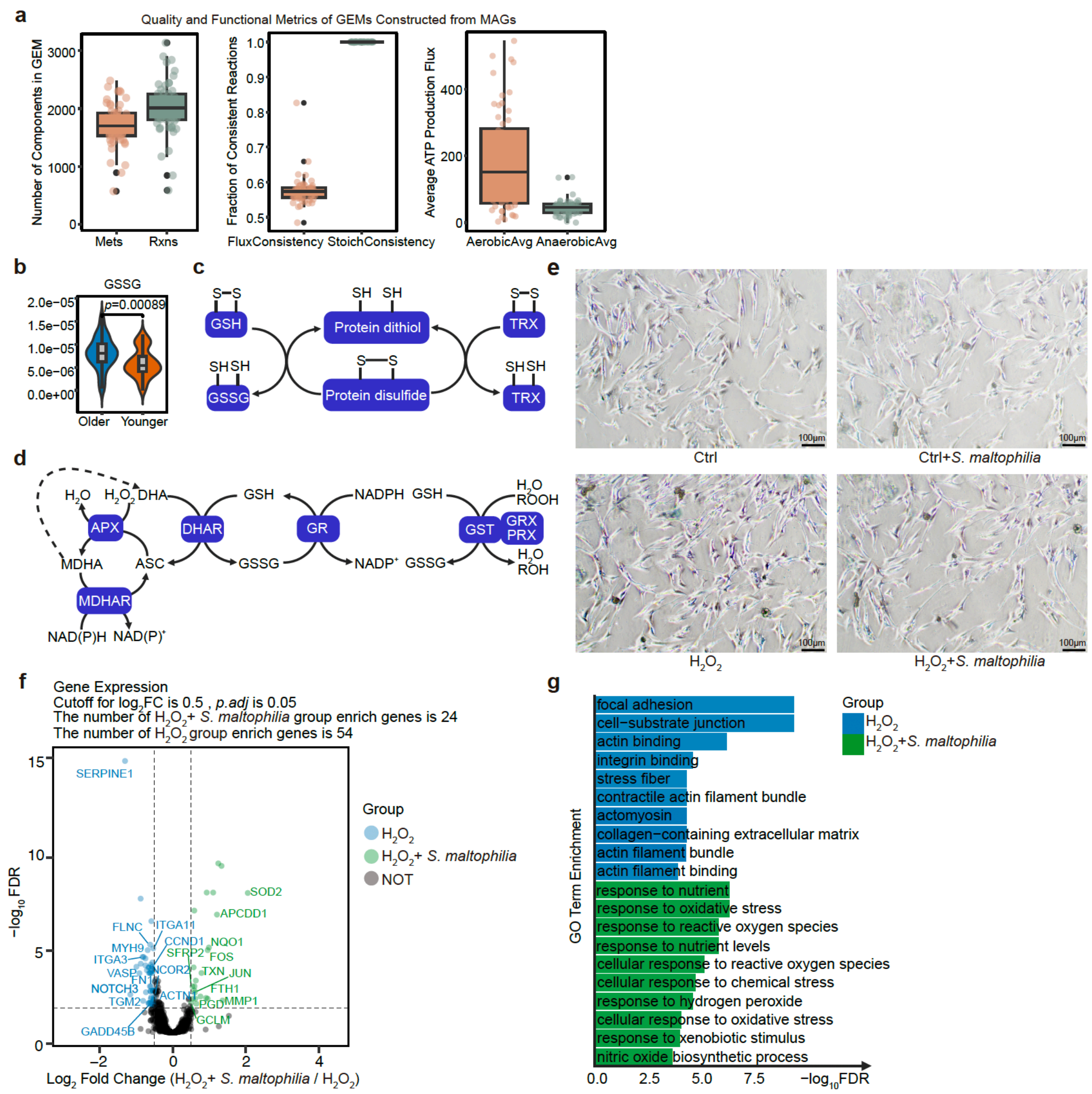


**Figure S4. Shadow price analysis from FBA illuminates antioxidant defense.**

(a) Reliability assessment of GEMs: Left Box Plot: Displays the number of reactions and metabolites for each MAG derived GEM. This visualization indicates the complexity and size of each model. Middle Box Plot: Shows the Flux Consistency and Stoichiometric Consistency for each GEM. These metrics evaluate the mathematical and biochemical robustness of the models, ensuring that they accurately represent the metabolic capabilities of the respective microbial communities. Right Box Plot: Compares the average ATP production under aerobic and anaerobic conditions across the GEMs.

(b) Violin plots depicting the significant differences in oxidized glutathione levels between older and younger groups in skin metabolomics. Notable differences are indicated, $P < 0.05$.

(c) This Figure demonstrates how glutathione (GSH) maintains proteins in their reduced thiol states within cellular environments. This reaction illustrates how GSH reduces protein disulfides to free thiol groups, thereby maintaining redox homeostasis and protein functionality.

(d) Glutathione-mediated antioxidant defense pathways: Left module: Ascorbate-dependent hydrogen peroxide metabolism: Monodehydroascorbate reductase (MDHAR) uses NAD(P)H to reduce monodehydroascorbate (MDHA) to ascorbate (ASC). Ascorbate peroxidase (APX) then uses ASC as an electron donor to convert $H_2O_2$ to water ($H_2O$). Middle module: Glutathione-dependent regeneration of ascorbate: Dehydroascorbate reductase (DHAR) reduces dehydroascorbate (DHA) back to ASC, consuming glutathione (GSH) and producing oxidized glutathione (GSSG). Right module: Glutathione-dependent peroxide metabolism: Glutathione reductase (GR) uses NADPH to convert GSSG back to GSH. Glutathione S-transferase (GST) and Glutaredoxin (GRX)/Peroxiredoxin (PRX) use GSH to reduce hydrogen peroxide ($H_2O_2$) and organic peroxides (ROOH) to water ($H_2O$) and alcohols (ROH).

(e) Representative images of skin primary fibroblasts exhibiting SA-β-Gal activity under different treatment conditions. Staining intensity indicates SA-β-Gal activity, with senescent fibroblasts appearing stained. Scale bars represent 100 μm.

(f) Volcano plot of differentially expressed genes between the $H_2O_2$ alone, and $H_2O_2$ treatment combined with *S. maltophilia* culture supernatant, analyzed using DESeq2. The analysis used an absolute $Log_2FoldChange > 0.5$ and adjusted $P < 0.05$.

(g) Bar plot shows the top 10 Gene Ontology (GO) terms with the smallest adjusted *P* enriched in the $H_2O_2$ alone, and $H_2O_2$ with *S. maltophilia* groups.

The GO terms include categories from Biological Process (BP), Molecular Function (MF), and Cellular Component (CC).

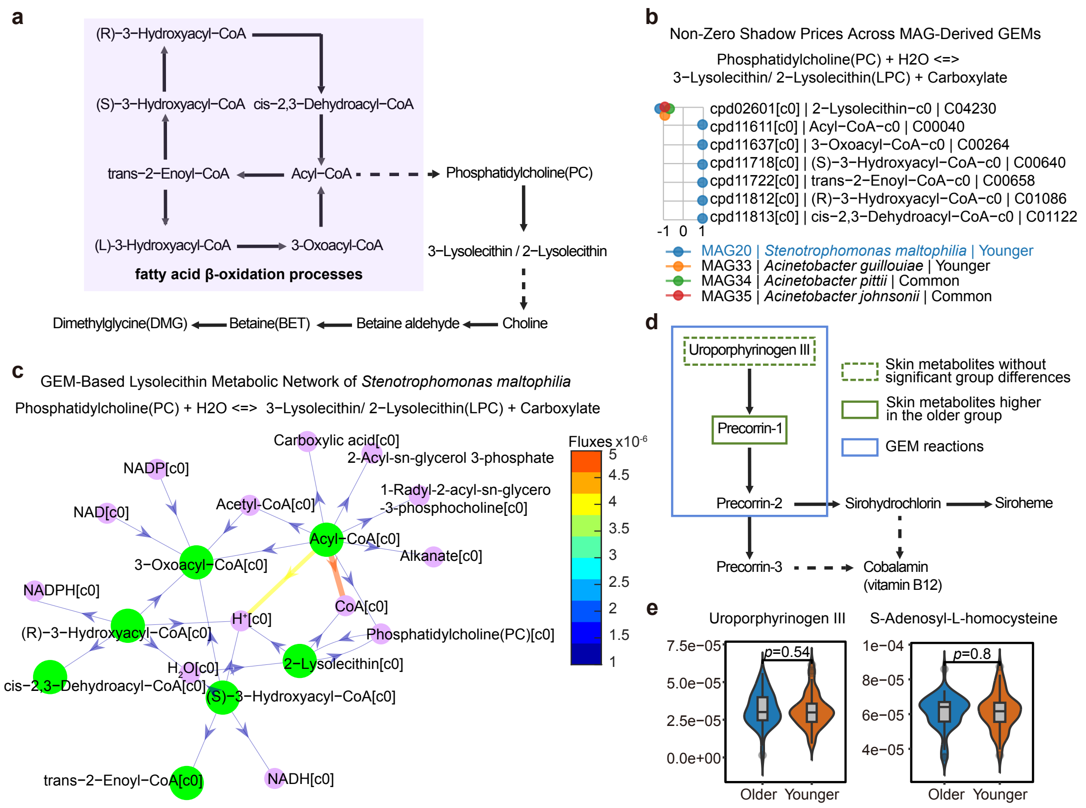


**Figure S5. Shadow price analysis from FBA illuminates betaine, lysolecithin synthesis dynamics, and porphyrin metabolism dynamics.**

(a) Schematic of the betaine metabolic pathway. This diagram illustrates the metabolic breakdown of lecithin/lysolecithin into choline, followed by subsequent conversions into betaine aldehyde, betaine (BET), and ultimately dimethylglycine (DMG) as a downstream product. Additionally, the fatty acid β-oxidation pathway is highlighted for its crucial role in generating acyl-CoA intermediates, which contribute to the synthesis and metabolism of lecithin/lysolecithin, thereby linking lipid metabolism with choline production. The dashed lines indicate simplified steps in the metabolic pathway, excluding intermediate metabolites for clarity.

(b) Shadow price analysis from FBA in lysolecithin synthesis reactions across different MAGs. The x-axis, y-axis, and color-coded dots follow a similar format to that in Figure 4a.

(c) Network and flux analysis of lysolecithin synthesis in *S. maltophilia* GEM: This provides a visual representation of the metabolic network involved in lysolecithin synthesis within *S. maltophilia*, annotated with flux values to indicate the intensity and direction of each metabolic step. The diagram serves to elucidate the complex interaction processes facilitating the synthesis of lysolecithin.

(d) Porphyrin and Vitamin B12 biosynthesis pathway in skin microbiome and host co-metabolism: This diagram illustrates the co-regulated metabolic pathway for porphyrin and Vitamin B12 synthesis involving skin microbiome and host interactions. The pathway initiates with Uroporphyrinogen III, leading to the sequential synthesis of Precorrin 1, Precorrin 2, and further downstream products, culminating in the production of Vitamin B12. Dashed green outlines denote skin metabolites with no significant differences between groups, solid green outlines indicate metabolites with higher abundance in the older group, and blue outlines represent GEM reactions.

(e) Violin plots illustrate the concentrations of critical metabolites, including Uroporphyrinogen III and S-Adenosyl-L-homocysteine, involved in the porphyrin metabolism pathway. The plots compare concentrations between older and younger groups. Despite variations, no statistically significant differences are indicated, $p$-value > 0.05.

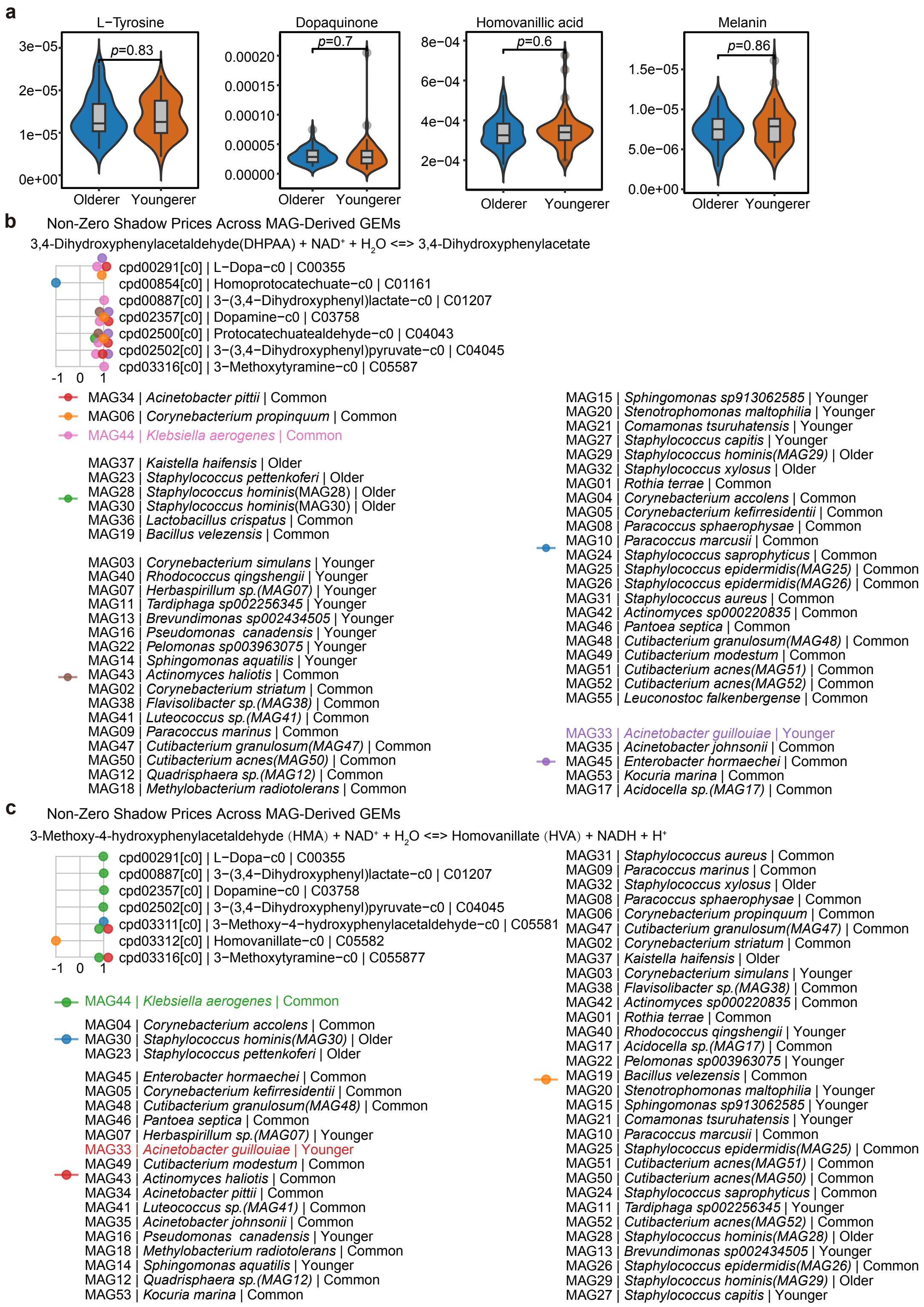


**Figure S6. Shadow price analysis from FBA illuminates tyrosine metabolism dynamics.**

(a) Violin plots illustrate the distribution of key metabolites involved in tyrosine metabolism within the skin metabolomics data. The plots compare concentrations between older and younger groups. Despite variations, no statistically significant differences are indicated, *p*-value > 0.05.

(b) FBA Shadow price analysis for the conversion of DHPAA to DOPAC in tyrosine metabolism: The Figure illustrates the shadow prices derived from FBA. The x-axis represents shadow price values, indicating the influence of each metabolite on this metabolic reaction. Positive values suggest that increasing the metabolite concentration drives the reaction forward. The colors represent different MAGs, highlighting the differential impact across microbial strains.

(c) FBA shadow price analysis for the conversion of HMA to HVA in tyrosine metabolism. The x-axis, y-axis, and color-coded dots follow a similar format to that in Figure S4B.

**Supplementary tables**

Table S1. Assessment of skin physiological traits, related to Figure 1.

Table S2. Metagenomic analysis of microbial species, diversity, and functional pathways, related to Figure 2.

Table S3. Comprehensive metabolomic analysis, differential core metabolites, and pathway enrichment, related to Figure 3.

Table S4. Analysis of metagenome-assembled genomes (MAGs) across samples, related to Figure S3.

Table S5. Genome-Scale Metabolic Modeling (GEM), experimental validation of *S. maltophilia'* s antioxidant and anti-aging roles, related to Figure 4.

Table S6. Comprehensive shadow price analysis of Betaine, Lysolecithin, Porphyrin, DHPAA, and HMA metabolism across MAGs in skin aging, related to Figures 5 and 6.
